# tRNA modifications are required for stress granule formation and melanoma metastasis

**DOI:** 10.1101/2025.08.20.671337

**Authors:** Riley O. Hughes, Hannah J. Davis, Leona A. Nease, Paul Zumbo, Mauro Danielli, Wyatt Tran, Kilian Maidhof, Kosmas Kosmas, Evgeny Kanshin, Shino Murakami, M. Graciela Cascio, Kellsey P. Church, Kelsey N. Aguirre, Ines Delclaux, Frank J. Slack, Ioannis S. Vlachos, Beatrix Ueberheide, Samie R. Jaffrey, Florian Erhard, Robert Banh, Doron Betel, Elena Piskounova

## Abstract

Metastasis is the leading cause of cancer related deaths, however therapies specifically targeting metastasis are lacking and remain a dire therapeutic need in the clinic. Metastasis is a highly inefficient process that is inhibited by extracellular stress. Therefore, metastasizing cells that ultimately survive and successfully colonize distant organs must undergo molecular rewiring to mitigate stress. Wobble uridine modifications, especially 5-methoxycarbonylmethyl-2-thiouridine (mcm^5^s^2^U_34_), have been implicated in stress response and poor prognosis of cancer patients. We use a patient derived xenograft (PDX) model of melanoma metastasis to study the role of the mcm^5^s^2^U_34_ modification in the stress response of metastasizing cells. We find that upon depletion of elongator acetyltransferase complex subunit 1 (ELP1)— a component of the mcm^5^s^2^U_34_ pathway on 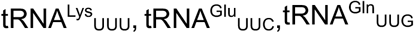, and 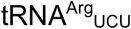 —codon-biased translation, migration, invasion, and metastatic burden *in vivo* is reduced. Further, we observe that stress granule components are enriched in a subset of codon-biased genes that are exclusively upregulated at the protein level in metastatic nodules compared to the primary tumor in our PDX model. Additionally, upon knockdown of ELP1, stress granule components have decreased protein expression with no significant change to their mRNA levels. Efficient translation, mediated by the carboxy-methylation arm of the mcm^5^s^2^U_34_ modification, is required for metastasizing cancer cells to withstand stress via stress granule formation and increase survival throughout the metastatic cascade. This makes the mcm^5^s^2^U_34_ machinery a potentially actionable therapeutic target, specific to metastatic disease.

## Introduction

Codon-biased translation has been established as a specific stress-resistance mechanism across many species.^1^ One of the molecular regulators of codon-biased translation is modification of the tRNA anticodon loop, including 5-methoxycarbonylmethyl 2-thiol on uridine 34 of tRNAs (mcm^5^s^2^U_34_).^1,2^ mcm^5^s^2^U_34_ has been shown to be essential for survival under stress in yeast and other organisms, but dispensable under non-stress conditions.^3-6^ Despite the demonstrated role of mcm^5^s^2^U_34_ in stress resistance in model organisms, the pathological role of mcm^5^s^2^U_34_ modifications remains poorly understood.

Modifications within the anticodon loop are critical for the interactions between the codon and anticodon, thereby regulating translational efficiency and fidelity.^3,5-8^ tRNA wobble uridine (U_34_) modifications have been shown to promote codon-biased translation in response to stress.^3^ U_34_ can be modified by a 5-carboxymethylation (cm^5^) mediated by the elongator acetyltransferase complex (ELP1-6) followed by a methylation (**m**cm5; bold indicates further 5-methylation) by alkylation repair homolog 8 (ALKBH8) (Figure 1A). A subset of mcm^5^U_34_ tRNA can also be modified with a 2-thiol group (s^2^) by the cytosolic thiouridylase complex (CTU1/2) on U_34_ (mcm^5^s^2^U_34_) of specific tRNAs (Figure 1A).^3,5,6^ However, other cm^5^U_34_-tRNAs are modified by different enzymes, resulting in **x**cm^5^U (bold × indicates other derivatives) modified tRNAs, including 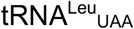 (Supplementary Figure 1A).^9^ Importantly, mcm^5^s^2^U_34_ has only been observed on the tRNAs that canonically pair with the A-ending codon in certain split codon boxes of the genetic code. Split codon boxes are unique in that the identity of the third base in the codon, which pairs with the wobble base of the anticodon (N_34_), is needed to determine the identity of the encoded amino acid (Supplementary Figure 1B). mcm^5^s^2^U_34_ exclusively occurs on 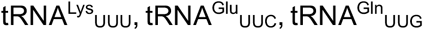, and 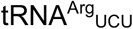. The Uridine-Adenosine base pair (U:A) is much weaker than the Guanosine-Cytosine pair and unmodified uridine is known to be promiscuous in its pairing, especially at the wobble base.^2,10,11^ In addition, it appears that N6-methyladenosine (m^6^A) modifications within mRNA codons further reduce the interaction of the U:A pair; the m^6^A modification on mRNA was recently characterized to promote mRNA degradation by slowing ribosomes during translation and causing ribosomal collision events.^12,13^ However, this slowdown at m^6^A sites could be alleviated in HEK293Ts by tRNAs harboring wobble base modifications, especially mcm^5^s^2^U_34_.^12,13^ Therefore, U_34_ modifications allow for increased translational efficiency and fidelity by restricting wobble pairing of uridine with non-cognate codons and simultaneously promoting true-cognate codon pairing.^3,4^ Despite this major role in translation and stress resistance, functional studies investigating the mcm^5^s^2^U_34_ modification in cancer progression are lacking.^14^

**Figure 1.**
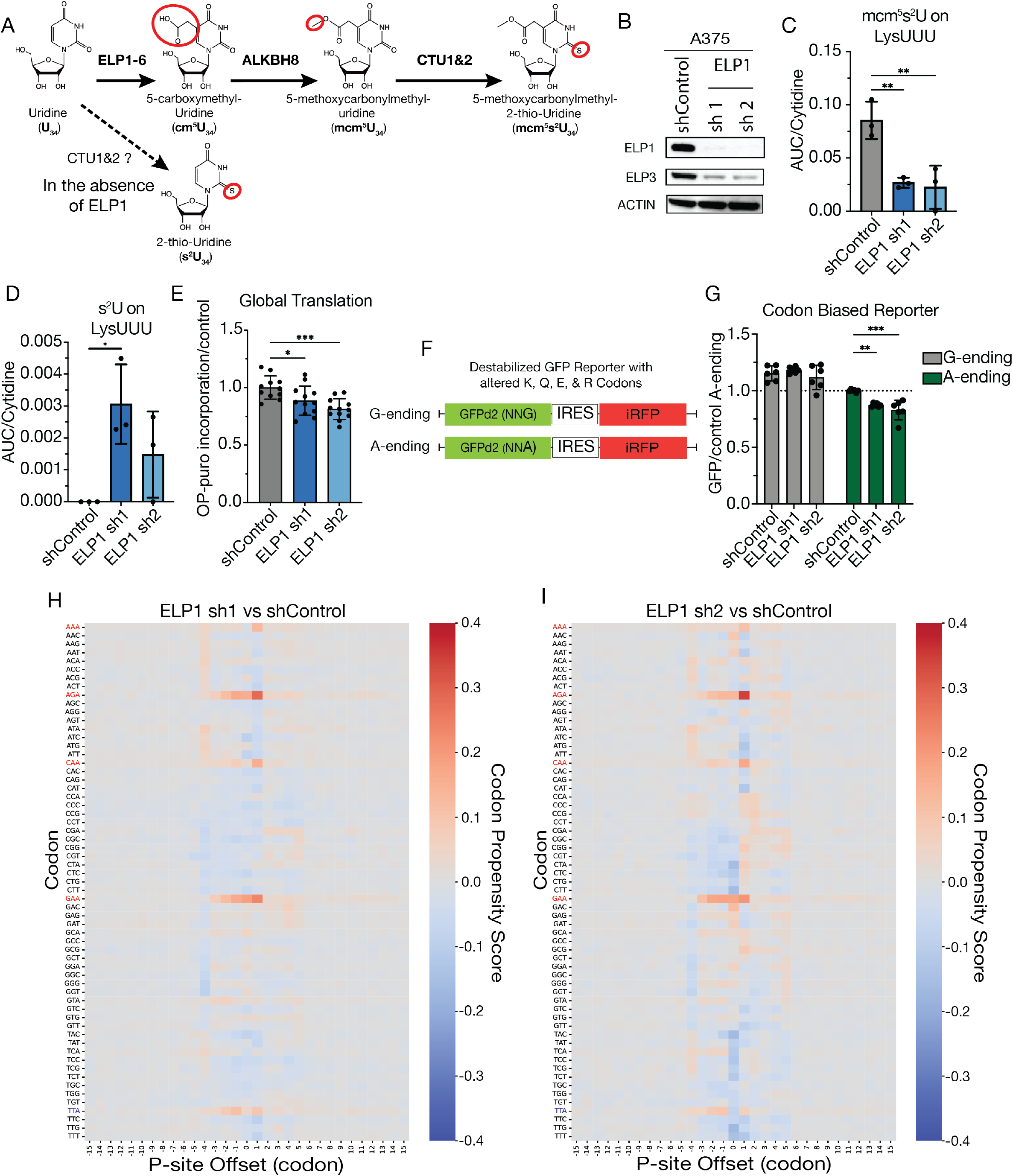
ELP1 loss reduces modifications of tRNA and impairs translation at specific codons. (A) The mcm^5^s^2^U_34_ pathway has been described to be a linear process starting with cm^5^ modification by ELP1-6, followed by further methylation at the 5-position (mcm^5^) by ALKBH8, and finally thiolation by CTU1&2. (B) Western blot of constitutive short hairpins targeting ELP1 in A375 melanoma cells. (C) Mean abundance (Area Under Curve, AUC) per cytidine (± SD) of mcm^5^s^2^ modified uridine on isolated 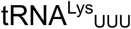 upon loss of ELP1 (blue) relative to control (gray). (D) The mean abundance (± SD) of s^2^ modified uridine on 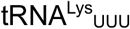 upon ELP1 loss relative to control. (E) Global translation measured by O-Propargal-puromycin (OP-puro) incorporation mean fluorescent intensity (± SD) normalized to control. (F) Schematic of the KQER (Lys, Gln, Glu, Arg) biased reporter utilizing a destabilized GFP (GFPd2) and IRES iRFP. (G) Loss of ELP1 with dox inducible hairpins reduces mean fluorescent intensity (± SD) of KQER GFPd2 with A-ending codons (green) but not G-ending codons (gray) measured by flow cytometry, normalized to signal from A-ending shControl. (H,I) Ribosomal footprint reads aligned to P-site with enrichment of codon propensity in the ELP1 knockdown lines shown in red and depletion of codon propensity shown in blue, mcm^5^s^2^U_34_ codons highlighted.

Targeting metastasizing cells has been challenging because of their intrinsic ability to adapt to extracellular stressors.^15,16^ Metastatic cells must undergo a series of rapid adaptations to survive under stress (e.g. oxidative, nutrient, hypoxia, etc.) as they transition through different environments and colonize vital organs.^15,17-22^ Such adaptability occurs by reprogramming the cellular proteome, often through transcriptional, epigenetic, or metabolic rewiring.^21,23^ There have yet to be any specific genetic alterations found that unambiguously endow cells with the ability to metastasize.^23,24^ Therefore, development of metastasis-specific therapies depends on the identification of specific reprogramming mechanisms that confer stress-resistance in cancer cells through mechanisms above the genome.

Stress granule assembly has been shown to regulate migration and invasion in various cancer types.^25,26^ Stress granules respond to various stress conditions, including those experienced by metastasizing cells, utilizing core RNA-binding proteins RAS GTPase-activating protein-binding protein 1 and 2 (G3BP1 and G3BP2) and Poly(A)-binding protein cytoplasmic 1 (PABPC1).^25-32^ Translationally stalled mRNAs can trigger stress granule formation by direct interaction with these RNA binding proteins leading to liquid-liquid phase separation of the RNA-protein granules. Further, Stress granule formation can allow for increased survival during stress by sequestration of mRNA transcripts, by acting as signaling hubs, and by promoting expression of other stress response proteins.^25,27,29^ For instance, increased stress granule formation in mutant KRAS-driven pancreatic cancer promotes drug resistance.^33^ However, it remains unclear how stress granule formation is regulated in metastatic cancers.

Limited studies have shown the impact of mcm^5^s^2^U_34_ modification in the stress response of human cancer. mcm^5^s^2^U_34_ was shown to play a role in resistance to targeted therapy in BRAF^V600E^ melanoma through regulation of HIF1α translation.^6^ Additionally, mcm^5^s^2^U_34_ promoted breast cancer tumor growth and progression through codon-biased translation of LEF1.^5^ However, the systemic role of mcm^5^s^2^U_34_ modifications in the context of metastasis has not been established. Recently, our group found that a single methylation event at the wobble uridine of the selenocysteine tRNA improved resistance to oxidative stress and was necessary for metastatic dissemination.^22^ Therefore we sought to further interrogate the role of the related wobble uridine modification, mcm^5^s^2^U_34_, in melanoma metastasis.

We hypothesized that the mcm^5^s^2^U_34_ modification regulates metastatic colonization by promoting translation of stress-response genes. We report that mcm^5^s^2^U_34_ modifications promote translation of codon-biased stress granule associated transcripts. Our work shows that loss of translational regulation upon ELP1 and mcm^5^s^2^U_34_ depletion reduces the ability of human melanoma cells to form stress granules and reduces cell motility *in vitro*. ELP1 loss also specifically reduces metastatic potential *in vivo*. Notably, we found that mcm^5^s^2^U_34_ is essential for stress granule formation by promoting the expression of core stress granule proteins exclusively through translational upregulation in metastases relative to primary tumors. This elongator-dependent loss of metastatic potential through the tRNA epitranscriptome and necessity for stress granule formation may represent an actionable therapeutic target for developing metastasis specific therapies.

## Results

### ELP1 knockdown reduces mcm^5^s^2^U_34_ levels and translation at specific codons

mcm^5^s^2^U_34_ is essential for stress resistance, a key feature for establishing successful metastases.^3,6,15,21,22^ Thus, we aimed to explore the role of 5-methoxycarbonylmethylation on U_34_ by ELP1 and ALKBH8. We targeted ELP1, the scaffold protein of the elongator complex, which is required for proper tRNA interaction with ELP3, the catalytic subunit.^34^ We observed that upon expression of either constitutive or doxycycline inducible short hairpins in human melanoma cells (A375 and A2058) against ELP1, the expression of ELP1 decreased (Figure 1B, Supplementary Figure 1C). We also observed a sympathetic decrease of ELP3 (Figure 1A and B). Further, analysis of 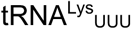 by mass spectrometry confirms a decrease in the mcm^5^s^2^U modification upon ELP1 knockdown (Figure 1C). This was accompanied by an increase in the presence of s^2^U modified nucleoside in the ELP1 knockdowns compared to control on the isolated 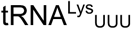 (Figure 1D). This indicates that the ELP1 knockdown is sufficient to reduce the deposition of the mcm^5^ mark on U_34_ and suggests that CTU1/2 modifies U_34_ without the modification at the 5-position.

Next, we assessed the impact of reduced of ELP1 on translation. Upon ELP1 knockdown, global translation was significantly decreased (Figure 1E). However, because the mcm^5^s^2^U_34_ modification occurs on 4 tRNAs (Lys, Gln, Glu, and Arg) and has been described to only improve translational efficiency of A-ending codons, we investigated if there were differences in the translation of these specific codons (AAA, CAA, GAA, and AGA) in A375 melanoma cells. We designed codon-biased genetic fluorescent reporters utilizing a destabilized GFP with a half-life of 2 hrs (GFPd2) with altered mcm^5^s^2^U_34_ dependent codon bias and a mcm^5^s^2^U_34_ independent iRFP for normalization (Figure 1F).^35^ Upon knockdown of ELP1 there was a decrease in translation of GFP encoded by A-ending codons at each occurrence of a Lys, Gln, Glu, or Arg (GFPd2 KQER A-ending), but did not observe any decrease in the reporter that was encoded by G-ending codons at the same amino acid positions (GFPd2 KQER G-ending) (Figure 1G). The observed decrease in expression of the A-ending GFP was approximately 20% suggesting that the decrease in global translation was mediated by decreased translation specifically at these 4 mcm^5^s^2^U_34_ dependent codons.

### Reduced mcm^5^s^2^U_34_ causes ribosomal stalling at specific codons

We next wanted to know if mcm^5^ on U_34_ of tRNA enhances the decoding rate of translating ribosomes. We performed ribosome profiling (Ribo-seq) in wildtype and ELP1 depleted A375 melanoma cells grown in either 2D adherent or 3D suspension culture, a general stress condition mimicking detachment and shown to induce oxidative stress (Supplementary Figure 2A). Here, we observe ribosome stalling when mcm^5^s^2^U_34_-dependent codons are in the A-site of ribosomes in A375 ELP1 knockdown cells compared to the control cells (Figure 1H and I, Supplementary Figure 1D and E). This A-site stalling is accompanied by an accumulation of ribosome footprint reads on mcm^5^s^2^U_34_-dependent codons (Lys^AAA^, Arg^AGA^, Glu^GAA^, and Gln^CAA^) upstream of the A-site, suggesting decreased efficiency of mcm^5^s^2^U_34_-dependent codon readthrough upon ELP1 depletion. Importantly, this includes all 4 mcm^5^s^2^U_34_-dependent codons including the Gln^CAA^ codon which is not known to receive m^6^A.^12^ Outside of the mcm^5^s^2^U_34_ dependent codons, there is an accumulation of footprints with the Leu^UUA^ codon upstream of the A-site that is less pronounced than the mcm^5^s^2^U_34_-dependent codons. This is likely due to the reduced carboxymethylation on the wobble uridine of 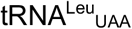 upon depletion of the elongator complex. These findings suggest that thiolation of U_34_ decreases translational efficiency when the nucleotide lacks mcm^5^. Taken together, this suggests that the decrease in global translation observed upon ELP1 loss is driven by the decreased translational efficiency of A-ending, mcm^5^s^2^U_34_ dependent, codons in endogenous mcm^5^s^2^U_34_ codon-biased transcripts.

### Reduced mcm^5^s^2^U_34_ decreases cell motility and blocks metastasis without affecting primary tumor growth

Next, we assessed the motility of melanoma cells with decreased mcm^5^s^2^U_34_ modification, as we predict loss of the modification will limit their metastatic potential. Melanoma cells with loss of ELP1 expression have significantly decreased capacity to migrate and invade *in vitro*, suggesting that this modification may be required for the metastatic potential of human melanoma *in vivo* (Figure 2A and B, Supplementary Figure 2B).

**Figure 2.**
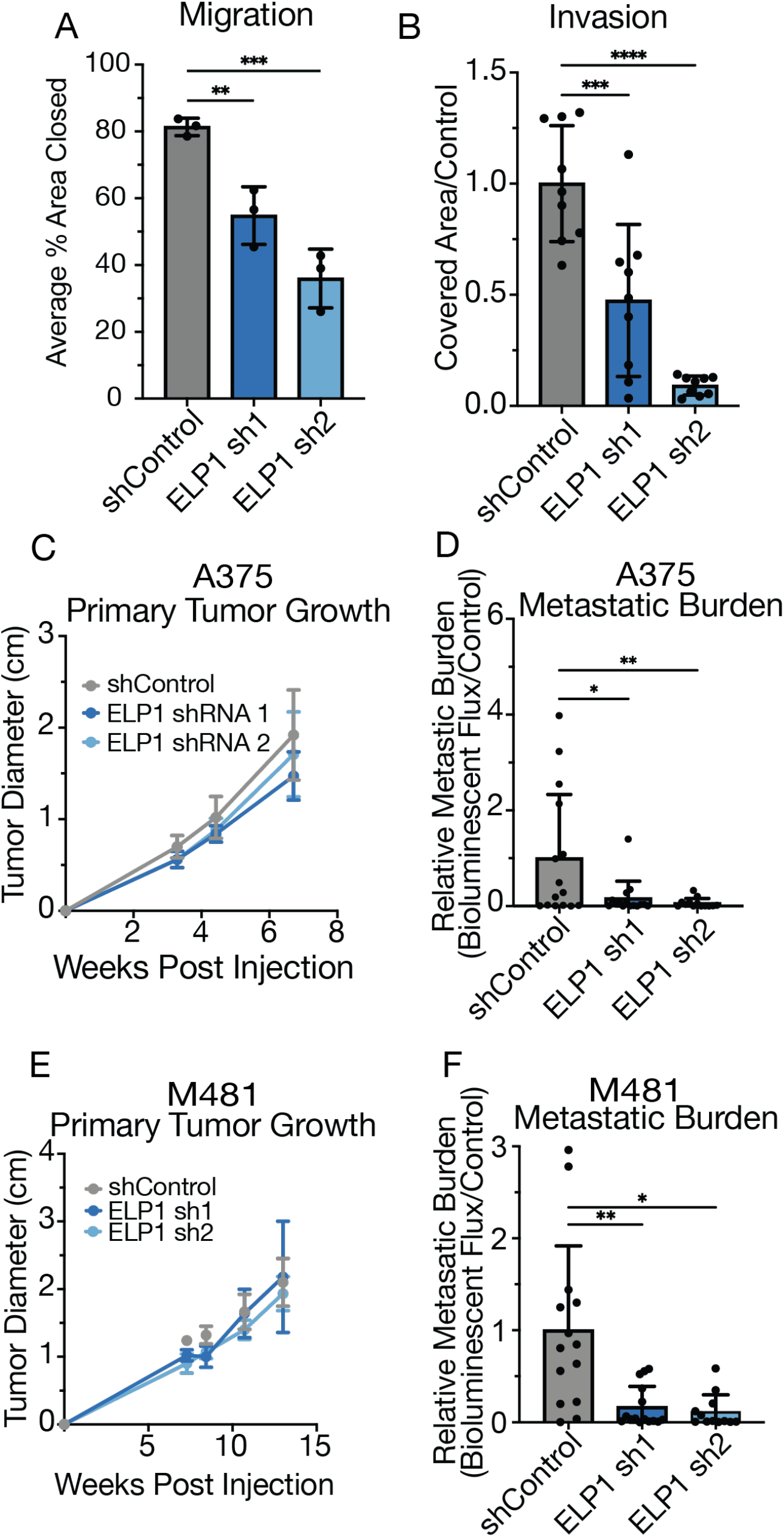
ELP1-mediated carboxymethylation of tRNA is necessary for metastatic potential. (A,B) Migration and invasion of A375 melanoma cells upon doxycycline inducible ELP1 loss (mean ± SD). (C,D) A375 melanoma cells orthotopically injected in NSG mice, shown is (C) primary tumor growth *in vivo* (D) and metastatic burden measured by bioluminescence signal (mean ± SD). (E,F) PDX melanoma, patient M481, orthotopically injected into NSG mice, shown is (E) primary tumor growth *in vivo* (F) and metastatic burden measured by bioluminescence signal (mean ± SD).

Therefore, we orthotopically injected ELP1 depleted A375 human melanoma cells into NOD.CB17-*Prkdc*^*scid*^ *Il2rg*^*tm1Wjl*^/SzJ (NSG) mice. We observed no change in primary tumor growth but a significant decrease in metastatic burden in NSG mice orthotopically injected with ELP1 depleted cells compared to control (Figure 2C and D). Likewise, patient derived xenograft (PDX) melanoma from two different patients, orthotopically injected into NSG mice, also showed decreased metastatic potential upon ELP1 loss with limited impact on primary tumor growth (Figure 2E and F, Supplementary Figure 2C-F). We observed the same phenomenon with ALKBH8 knockdowns (Supplementary Figure 2G-I), but as the ALKBH8 mediated methylation event to produce mcm^5^U_34_ from cm^5^U_34_ cannot occur without the prior action of the elongator complex (to first generate cm^5^U_34_) we focused on ELP1 depletion (Figure 1A). Our data implicates the necessity of ELP1 and ALKBH8, the mcm^5^ arm of the mcm^5^s^2^U_34_ modification pathway, in metastatic potential *in vivo* and *in vitro*. This modification is associated with translational regulation and stress resistance, so we next aimed to uncover the role of ELP1 in translation in human melanoma metastasis.

### Codon-biased translation supports cell motility

To assess codon biased translation *in vitro*, we expressed the codon biased reporters in melanoma cells, with no alteration to ELP1 expression. We observed that melanoma cells (A375 and A2058) that efficiently expressed the A-ending version of the KQER reporter were significantly more motile than those cells that inefficiently expressed the reporter. In comparison, there was no difference in motility between the cells efficiently or inefficiently expressing the G-ending reporter or a non-destabilized GFP overexpression control (Figure 3A-F). In effect, migratory capacity can be predicted by the ability of cells to efficiently express protein products of transcripts biased towards usage of mcm^5^s^2^U_34_ dependent codons (Figure 3C and F). This suggests that codon biased translation supports cell motility phenotypes.

**Figure 3.**
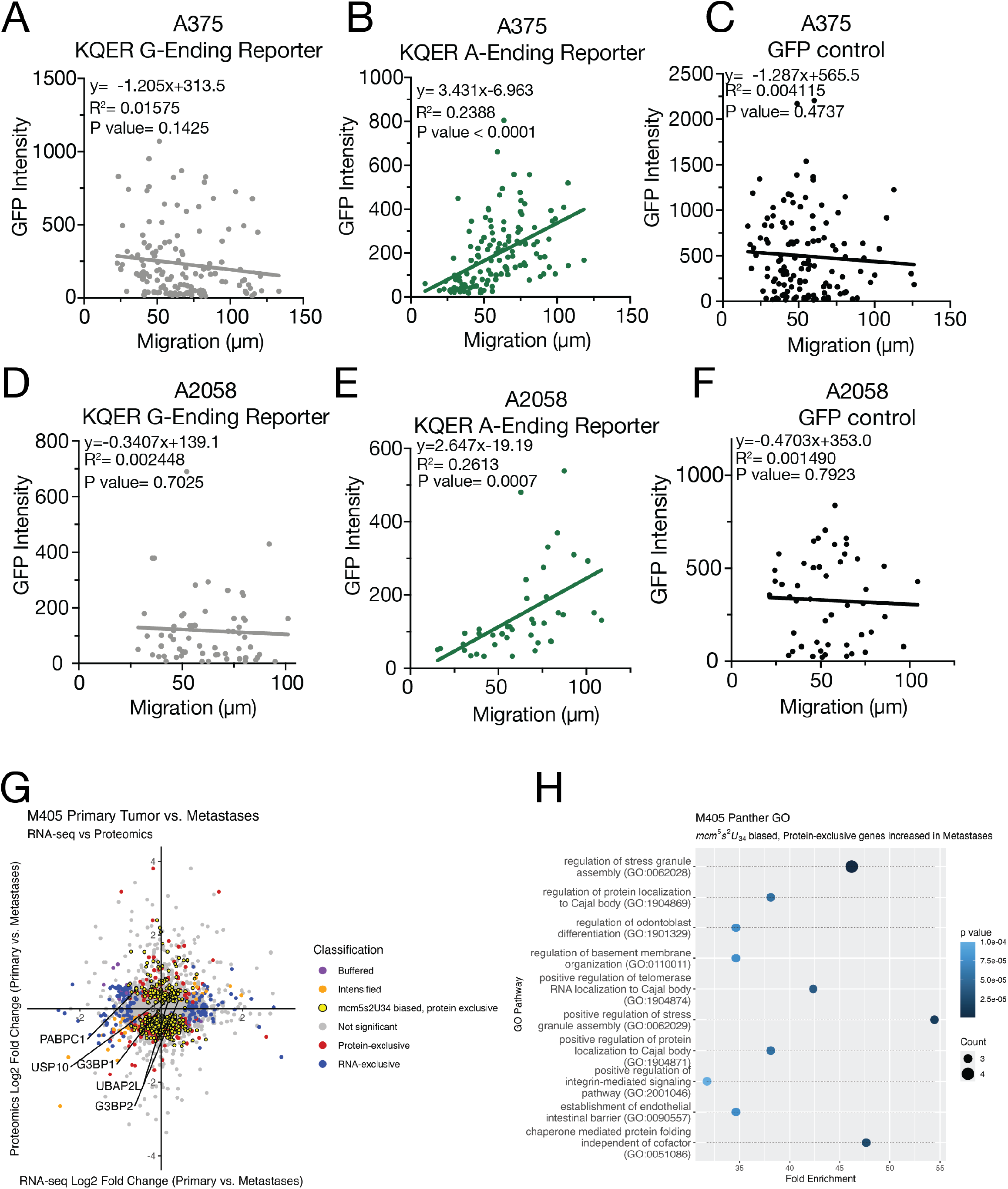
Metastatic phenotypes are associated with increased codon-biased translation of mcm^5^s^2^U_34_ decoded stress response mRNAs. (A-F) Correlation of codon-biased reporter expression with single-cell migration. (A-C) Scatter plot of total migration distance vs. GFP intensity of A375 melanoma cells expressing either the KQER GFPd2 (A) G-ending or (B) A-ending reporters or (C) a control, non-destabilized GFP overexpression construct. Solid lines indicate least squares regression fit (*y=mx+b*). *m, b*, R^2^, and two-tailed Pearson P value are shown. (D-F) Scatter plot of total migration distance versus GFP intensity of A2058 melanoma cells expressing either the KQER GFPd2 (D) G-ending or (E) A-ending reporters or (F) a control, non-destabilized GFP overexpression construct. Solid lines indicate least squares regression fit (*y=mx+b*). *m, b*, R^2^, and two-tailed Pearson P value are shown. (G-H) Multiomics plot of log2 fold change of both RNA-seq and Proteomics data in metastases vs primary tumor of samples collected from a PDX model, patient M405, of melanoma metastasis, with associated gene ontology terms for the set of genes that were exclusively increased at the protein level and had codon biases for A-ending mcm^5^s^2^U_34_ codons.

Progression through the metastatic cascade requires sufficient management of external stress.^15^ To investigate specific transcripts that are impacted by loss of ELP1 under stress conditions *in vitro*, we examined translational efficiency (TE) by quantifying ribosome footprints on each transcript relative to the expression of the transcript (Supplementary Figure 3A-D). Of note, the RNA binding proteins PABPC1, PABPC3, and PABPC4— known to be involved in stress granule assembly and maintenance— all had significantly decreased TE in ELP1 sh2 under stress-inducing 3D growth conditions, but not in the 2D conditions. These RNA binding proteins are known to be involved in stress granule assembly and maintenance, suggesting these may be transcripts that are necessary for survival especially under stress (Supplementary Figure 3A-D).

Taken together, this suggests that the tRNA modification state is necessary for codon biased translation that support stress resistance and contributes to cell motility events. Further this codon-biased translational decrease across stress transcripts upon ELP1 loss may be what is driving the decrease in metastatic potential observed *in vivo*. Due to the translational phenotype of ELP1 loss *in vitro* and the metastatic phenotype of ELP1 loss *in vivo*, we aimed to investigate the role of translational regulation in metastasis by identifying endogenous mcm^5^s^2^U_34_ codon-biased transcripts that may be translationally upregulated in metastasis.

### Core Stress Granule Components are associated with codon biased translation and metastasis

We identified codon-biased genes solely changing at the proteomic level and not the RNA level in metastases relative to primary tumor. Due to the technical challenge of doing Ribo-seq in tissue, we took a multiomics approach and analyzed RNA-seq and proteomics data taken from our *in vivo* PDX models of metastasis, defining sets of genes that were exclusively changing in either the transcriptome or the proteome (Figure 3G, Supplementary Figure 3E). We also defined genes as having their protein expression either buffered (significant log2 fold changes of RNA-seq vs. proteomics in opposite directions) or intensified (significant changes in the same direction) by the mRNA expressioin.^36^ We reasoned that the subset of genes for which the change in expression between metastases and primary tumor at the level of protein is significantly different, but is not explained by changes at the transcript level, could be candidates for translational regulation in metastatic disease. However, these genes may have no change in transcript expression due to any post-transcriptional regulatory processes, including regulation of translation or post-translational processing.

Next, we wanted to know if elongator mediated mcm^5^ regulates translation of metastatic protein-exclusive proteins. We first defined transcripts that may be regulated by the mcm^5^s^2^U_34_ modification (which occurs exclusively on 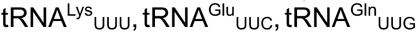, and 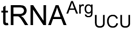) by examining their codon bias using relative synonymous codon usage (RSCU) score (Equation 1).^37^ Using this RSCU score, we further defined a subset of the protein exclusive genes that had a codon bias towards usage of mcm^5^s^2^U_34_ dependent codons. We preformed gene ontology pathway analysis on this subset of mcm^5^s^2^U_34_ codon-biased, protein-exclusive genes with positive changes in metastatic lesions relative to primary tumor of two different PDX melanomas (Figure 3H, Supplementary Figure 3F). We observed enrichment in pathways such as protein folding, protein transport, and notably stress granule assembly. It has been recognized that metastasizing cells must overcome various forms of external stress to successfully colonize distant organs, therefore we were particularly intrigued by enrichment of the stress granule pathway in this dataset. The stress granule associated genes we observe in this subset of genes includes: G3BP1, G3BP2, UBAP2L, USP10, and PABPC1. Notably G3BP1 and G3BP2, and PABPC1 are core components of stress granule assembly.^30,32,38^ Since we also observed PABPC1 translation to be affected *in vitro* in 3D conditions, we investigated if mcm^5^ promotes stress granule formation more directly.

### Stress Granule formation is decreased upon loss of ELP1

Intrigued by the enrichment of the stress granule assembly pathway in the codon-biased metastatic proteomes of PDX melanoma, we assessed the ability of ELP1 KD cells to form stress granules induced by treatment with sodium arsenite (NaAsO_2_). Immunofluorescent staining of G3BP1 and PABPC1 both show significantly decreased percentage of cells able to form stress granules in ELP1 knockdown compared to control (Figure 4A-D).

**Figure 4.**
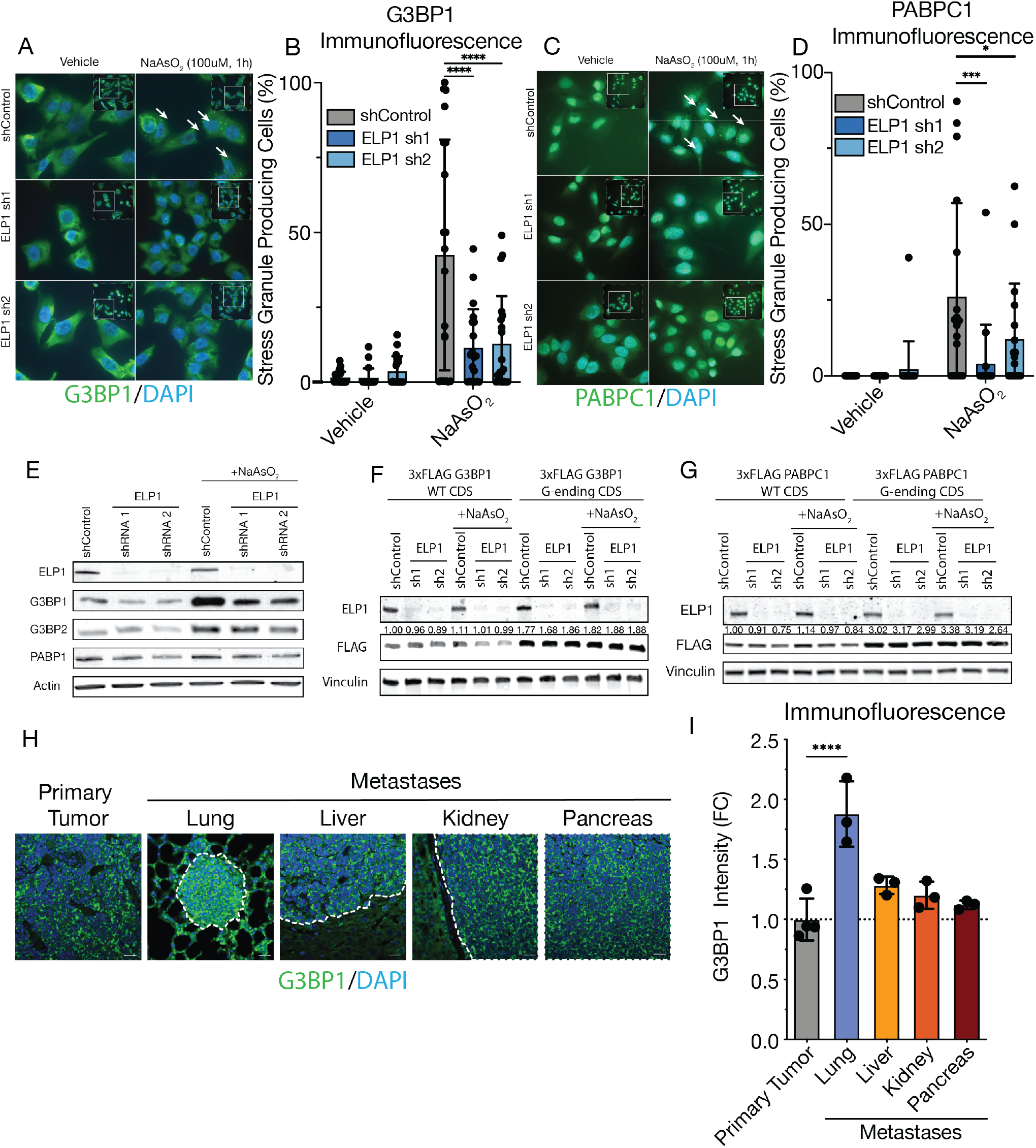
Stress granule components decrease upon ELP1 KD in a mcm^5^s^2^U_34_ codon-biased dependent manner and are elevated in metastases. (A-D) Immunofluorescent staining of A375 melanoma cells treated with 100 μM AsNaO_2_ for 1 hr. (A, B) Representative images and quantification of G3BP1 staining (mean ± SD). (C, D) Representative images and quantification of PABPC1 staining (mean ± SD). (E-G) Western blot for of A375 melanoma cells treated with 100 μM AsNaO_2_ for 1 hr. (E) Expression of Stress granule components upon loss of ELP1. (F, G) Representative images of overexpression of either wild type (WT) coding sequences (CDS) or G-ending codon switched CDS of stress granule components, (F) G3BP1 and (G) PABPC1, upon loss of ELP1. Values indicate mean FLAG signal from 3 independent experiments relative to vinculin control and normalized to untreated control. (H-I) Immunofluorescent imaging of tissue sections taken from mice injected with M481 PDX melanoma, (H) shown is the primary tumor and metastatic nodules (outlined in dashed white line) in various colonized organs with DAPI and G3BP1 staining and the (I) quantification of immunofluorescent signal normalized to primary tumor (mean ± SD).

We next wanted to know if the reduced stress granule formation in ELP1 cell after arsentie treatment is due to reduced translation of mcm^5^s^2^U_34_ decoded codon-enriched transcripts such as G3BP1, G3BP2, and PABPC1. ELP1 knockdown cells treated with NaAsO_2_ had decreased expression of the stress granule core proteins G3BP1, G3BP2, and PABP1 (Figure 4E). This reduced expression of the core proteins is likely due to reduced translation since their mRNA expression is either not significantly changing or increasing upon ELP1 loss and NaAsO_2_ treatment (Supplementary Figure 4A-C). Taken together, these results indicate that ELP1 expression and the associated mcm^5^ modification are required for expression of stress granule core components and stress granule formation under stress. Further, this suggests that depletion of ELP1 and the associated codon-specific reduction in translational efficiency makes cells unable to adequately respond to stress.

Next, we wanted to directly test whether the reduced translation of G3BP1, G3BP2, and PABPC1. Is due to the codons decoded by mcm^5^s^2^U_34_-modified tRNAs. To do so we generated G3BP1, G3BP2, and PABPC1 reporters, much like the reporters in Figure 1F. these reporters were inserted into overexpression plasmids with a 3xFLAG tagged G3BP1, G3BP2, or PABPC1 coding sequence (CDS) with either the wild type codons or altered all occurrences of A-ending mcm^5^s^2^U_34_ codons to their synonymous G-ending counterparts. We observed that expression of the WT CDS, was decreased upon ELP1 KD but ELP1 loss had no impact on the expression of the G-ending versions of the CDS (Figure 4F and G, Supplementary Figure 4D). Further, we assessed the expression of G3BP1 in the primary tumor and metastatic nodules in lung, liver, kidney, and pancreas in our PDX model of melanoma by immunofluorescent staining. G3BP1 expression was significantly elevated in melanoma metastases in the lung and trended towards increased expression in metastatic nodules that landed in other organs, relative to their matched primary tumors (Figure 4H-I).

This data, taken together with our observation that ELP1 loss reduces the ability of cells to metastasize, shows that stress granule assembly is a downstream target of decreased translational efficiency upon ELP1 loss and efficient translation of stress granule components may be required for metastatic progression.

## Discussion

It has been shown that only about 40% of variation in protein can be explained by changes in RNA expression, whereas the remaining 40-50% and 10-20% can be explained by changes to translational efficiency and protein degradation respectively.^39,40^ These findings point out that translational regulation is a major step in regulating gene expression. Emerging studies also suggest that translation can be regulated at multiple steps, many of which involve tRNA modifiactions. Recent work in HEK293Ts, MEFs, and mESCs has shown the importance of m^6^A on mRNAs in the regulation of ribosome dynamics and mRNA degradation, and that the efficient decoding of m6A-containg codons is mediated by mcm^5^s^2^U_34_ containing tRNAs.^12,13^ It was suggested that the relative expression of m^6^A writers vs mcm^5^s^2^U_34_ writers could predict outcomes and hazard ratios in patients across a wide spectrum of cancer types.^12^ Previous work in BRAF-therapy resistant melanoma identified HIF1α as a specific factor regulated by mcm^5^s^2^U_34_ at the level of translation.^6^ mcm^5^s^2^U_34_ modifications have also been shown to play a role in breast cancer tumor growth and progression through translational regulation of LEF1.^5^ In our study, we observed a more unique scenario, where loss of the mcm^5^ at U_34_ has no effect on the growth of the primary tumor in three different orthotopic models of melanoma but specifically inhibits metastatic colonization. Intriguingly, we do not see changes in either LEF1 or HIF1α in our melanoma cells *in vitro* or *in vivo* upon loss of mcm^5^s^2^U_34_ (Figure 3C, Supplementary figure 3C-G). This suggests that the function of mcm^5^s^2^U_34_ is highly context dependent; the modification may differ between cancer types and throughout steps of tumor progression. This invokes the likelihood of additional mechanisms of regulation of codon-biased translation that work in concert with mcm^5^s^2^U_34_ throughout these processes.

Here we show that ELP1, the scaffold protein of the cm^5^U_34_ writer of the mcm^5^s^2^U_34_ modification on tRNAs, is required for melanoma cell migration and invasion *in vitro*, and is specifically required for metastatic potential in melanoma PDXs *in vivo*, while being dispensable for the growth of the primary tumor. Further, efficient expression of a mcm^5^s^2^U_34_ dependent codon-biased reporter (GFPd2 KQER A-ending) could predict migratory capacity of melanoma cells. In fact, the A-ending mcm^5^s^2^U_34_ codon-biased proteins that are elevated in metastases of PDXs, specifically G3BP1, G3BP2, and PABPC1, are core components of a stress response pathway that has been shown to increase fitness of cancer cells.^33^ We show that G3BP1, G3BP2, and PABPC1 expression as well as stress granule formation under stress is decreased by loss of the mcm^5^s^2^U_34_ modification. Additionally, in a similar fashion as the KQER GFPd2 A-ending reporter, we show G3BP1, G3BP2, and PABPC1 have no response to ELP1 loss when their codon bias is altered to contain only G-ending synonymous codons. This codon specific translational impact is confirmed by transcriptome-wide ribosome footprinting. We see an accumulation of reads occur that have either a mcm^5^s^2^U_34_ dependent codon in the ribosomal A-site or recently in the ribosomal A-site in ELP1 knockdown cells indicating ribosomal stalling upon encountering these specific codons (AAA, CAA, GAA, and AGA). Importantly this includes the CAA codon which does not receive the m^6^A modification; this observed accumulation of reads with the CAA codon at or around the A site in ELP1 knockdowns relative to control may suggest cancer specific ribosomal dynamics as this accumulation pattern was not observed in HEK293Ts (Figure 3A-B), and is likely not dependent on the presence of m^6^A.^12^

Still, these findings suggest that metastasizing tumor cells rely on efficient translation of these core stress granule components, potentially to deal with metastatic stress. Stress granules have previously been implicated in the chemoresistance of pancreatic cancer through a mutant KRAS dependent manner, though the ubiquitousness of this mechanism warrants further investigation.^25,33,41^ Additionally, stress granule components themselves have been associated with migration, invasion, and metastatic phenotypes in various cancer types and their high expression can be a poor indicator of patient outcomes.^25,27^ This is fitting as cancers in these later stages of progression, like drug resistance and metastasis, must overcome external stressors to survive. As a response to stress, the IDR2 domain of G3BP1 interacts with protein kinase R (PKR) of the integrated stress response (ISR) and drives phosphorylation of eukaryotic elongation factor 2α and stress granule nucleation.^27^ Further, G3BP1, G3BP2, and PABPC1 are components of a stress response in which cessation of stress can result in the autophagic turnover of the proteins involved in stress granule formation.^28,29^ In this way, core components of stress granules may be used for stress clearance and then degraded once the stress has been managed. Therefore, proteinaceous components of stress granules must be translationally replaced *de novo* to respond to the next onslaught of external stress, unlike stress response enzymes that can cyclically perform their stress clearance functions. This indicates that translational efficiency of A-ending codons in the metastatic setting is maintained by mcm^5^s^2^U_34_ modifications to allow a suitable amount of translation for survival. Additionally, the protein products of mcm^5^s^2^U_34_ dependent codon biased transcripts may be preferentially degraded more, potentially to deal with sustained or multiple bouts of cellular stress—requiring mcm^5^s^2^U_34_ modifications to maintain efficient translation for their *de novo* translation. Epitranscriptomic modification of tRNAs by mcm^5^s^2^U_34_ therefore represents a strategy for cells to continuously sustain efficient translation and replenishment of crucial proteins required for responding to stress throughout metastasis, emphasizing their necessity for metastatic progression. Therefore, perturbing the elongator complex, the cm^5^U_34_ epitranscriptomic writer in the mcm^5^s^2^U_34_ tRNA modification pathway, offers an exciting therapeutic opportunity for specifically targeting the process of metastasis in melanoma.

### Limitations of the Study

Previous work had identified the mcm^5^s^2^U_34_ pathway in regulating translational efficiency and implicated it in cancer progression. In this study, we characterized the impact of this pathway on translation in melanoma cells *in vitro*, but had limited success directly quantifying translation *in vivo*. Quantifying translation in metastatic nodules of *in vivo* models of melanoma remains a challenge due to current limitations both in resolving the contribution of contaminating mouse tissues in Ribo-seq and in abundance of starting material. Similarly, direct measurement of mcm^5^s^2^U_34_ modification levels is limited by low starting material and low throughput of the tRNA isolation method both from *in vivo* models of melanoma as well as from patient samples. Future efforts would benefit from directly measuring changes to this pathway in both modification status and protein expression in patient samples to validate this pathway as a viable therapeutic target for metastatic disease. Additionally, advances in ribosome profiling techniques for application *in vivo* to faithfully identify human ribosome protected fragments while limiting contamination of ribosome protected fragments derived from mouse tissues would improve our ability to identify *bona fide* translational targets in complex models of human disease.

## Acknowledgments

The authors would like to thank Lukas Dow for contributing the mir-E plasmids that were modified in this study, Maider A. Amiama at the Weill Cornell Medicine Applied Bioinformatics Core for her guidance on preliminary sequencing analysis, Nayah Bullen for her assistance with mouse processing, John Blenis for his guidance and mentorship, and the Beth Israel Deaconess Medical Center - BIDMC-Immunostaining and Microscopy Core, RRID:SCR_012312 for their imaging capabilities. 1T32GM141949 to R.O.H. Pershing Square Sohn Prize Cancer Research Prize to E.P. NCI Grants R37CA289040 to R.S.B.

---

**Supplementary Figure 1.**
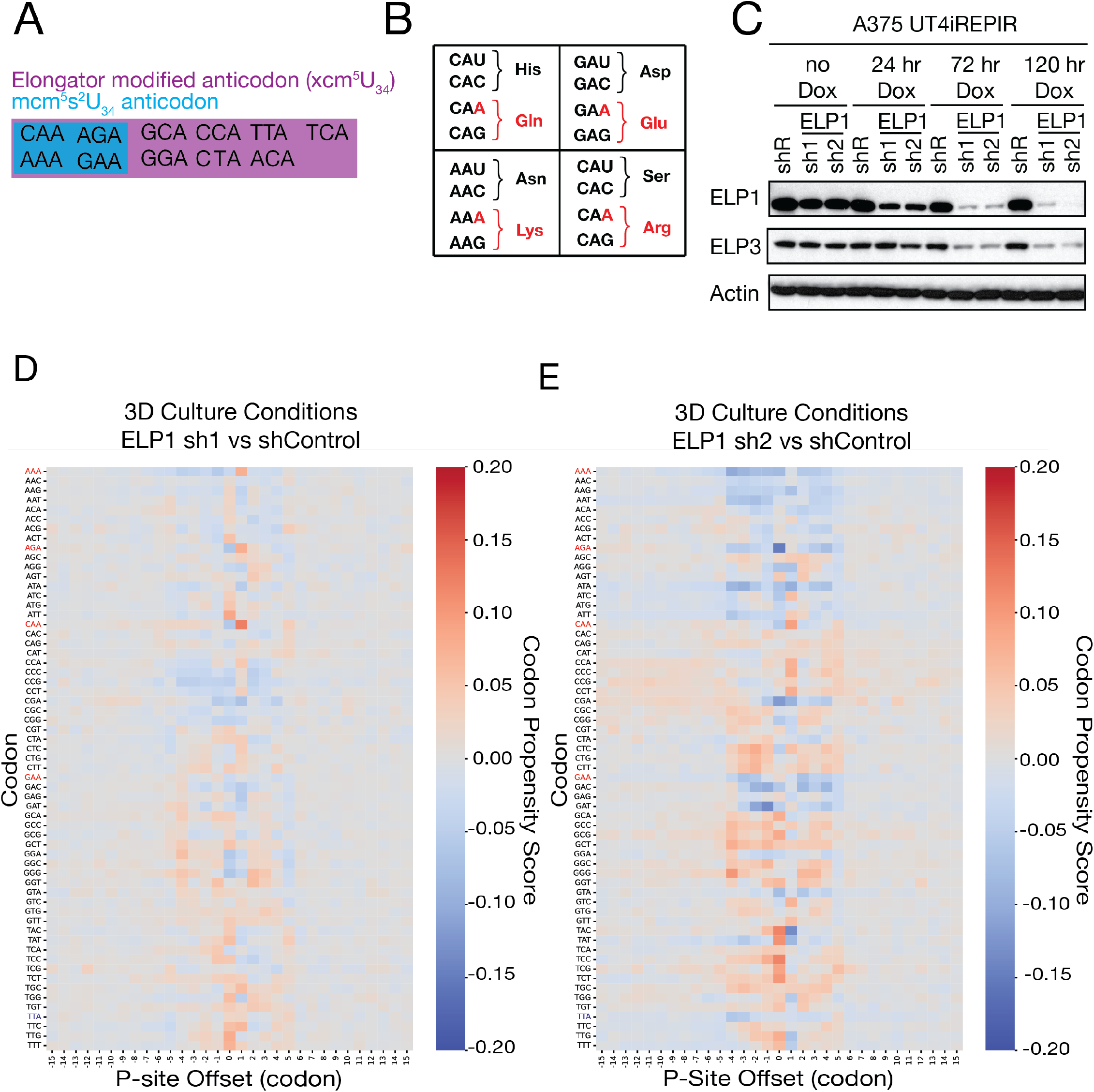
(A) Codons that receive the xcm^5^U_34_ modification by the elongator complex and the subset of those codons that receive the mcm^5^s^2^U^34^ version of the modification. (B) The genetic code contains split codon boxes, the indicated (Red) A-ending codons pair with mcm^5^s^2^U_34_ modified tRNAs, whereas the G-ending synonymous codons do not. (C) Doxycycline (dox) inducible short hairpins against ELP1 over a dox time course. (D-E) Ribosomal footprint reads from A375 ELP1 knockdown cells grown in 3D culture conditions for 24 hrs aligned to P-site with enrichment of codon propensity shown in red and depletion of codon propensity shown in blue, mcm^5^s^2^U_34_ codons highlighted.

**Supplementary Figure 2.**
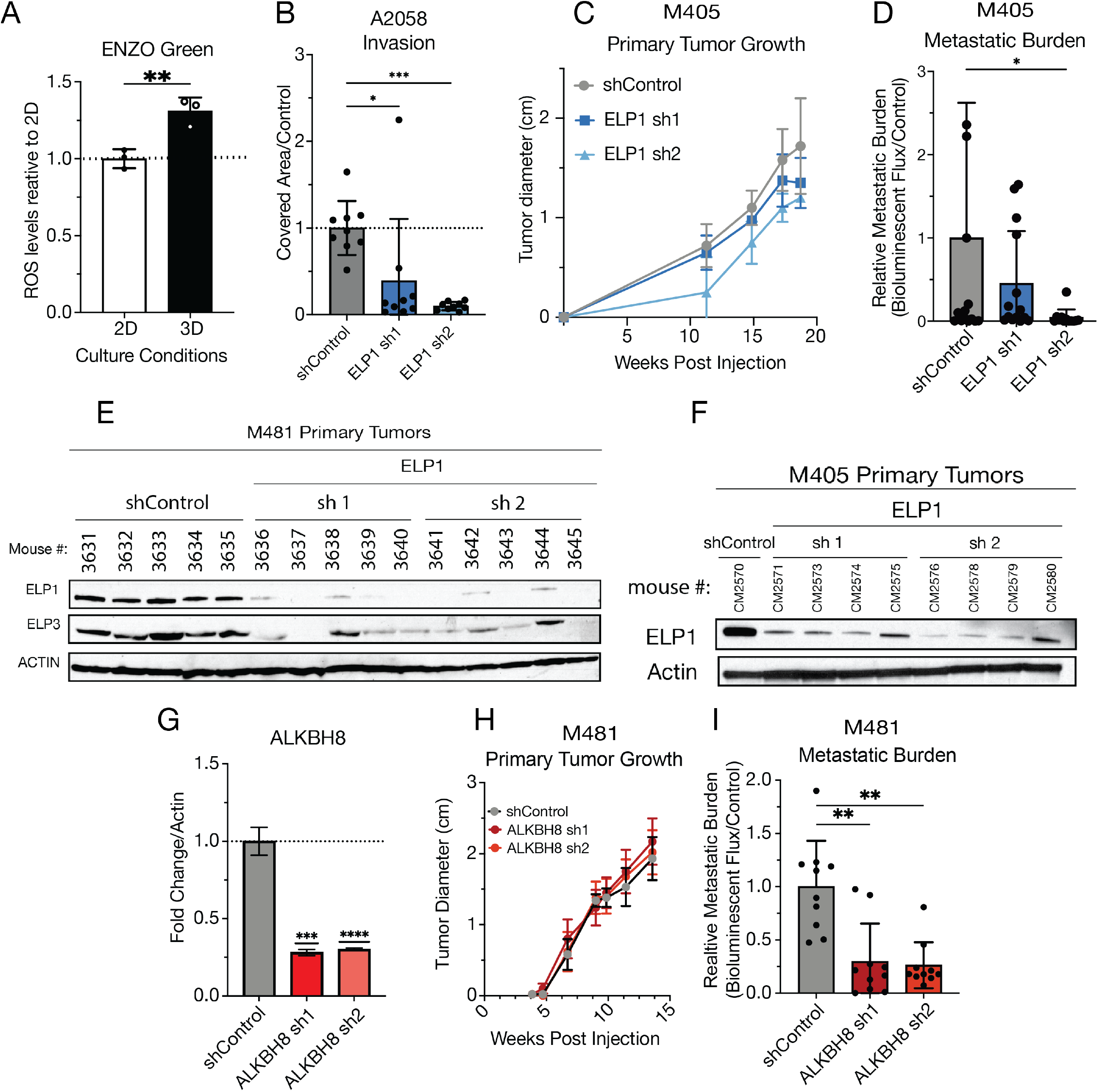
(A) Reactive oxygen species (ROS) measured by ENZO Green ROS-reactive fluorescent dye and flow cytometry in A375 melanoma cells cultured in 2D or 3D conditions. (B) Migration of A2058 melanoma cells is significantly reduced upon ELP1 loss. (C-D) PDX melanoma, patient M405, orthotopically injected into NSG mice, shown is (C) primary tumor growth *in vivo* (D) and metastatic burden measured by bioluminescence signal (mean ± SD). (E, F) Expression of ELP1 in primary tumors of mice at experimental endpoint for M481 and M405 PDX melanomas. (G-I) M481 PDX melanoma expressing ALKBH8 short hairpins orthotopically injected into NSG mice, shown is (G) Hairpin validation by qPCR (H) PDX primary tumor growth *in vivo* (I) and metastatic burden measured by bioluminescence signal (mean ± SD).

**Supplementary Figure 3.**
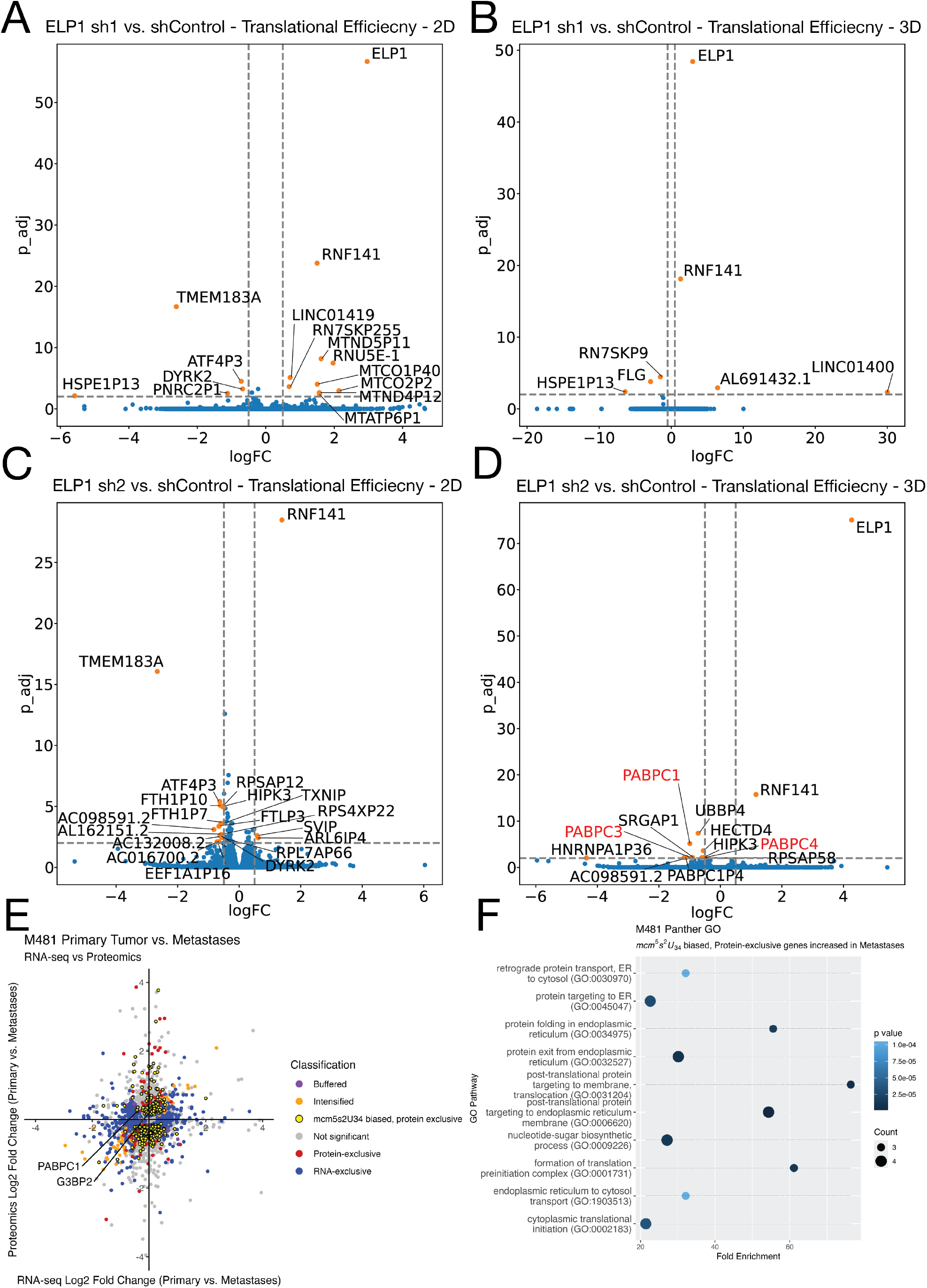
(A,B) Translational efficiency (TE) analysis of ribosome footprint reads in 2D and 3D culture conditions, notably PABPCs have a decrease in TE in ELP1 sh2 grown in 3D conditions. (G,H) Multiomics plot of log2 fold change of both RNA-seq and Proteomics data in metastases vs primary tumor of samples collected from a PDX model, patient M481, of melanoma metastasis, with associated gene ontology terms for the set of genes that were exclusively increased at the protein level and had codon biases for A-ending mcm^5^s^2^U_34_ codons.

**Supplementary Figure 4.**
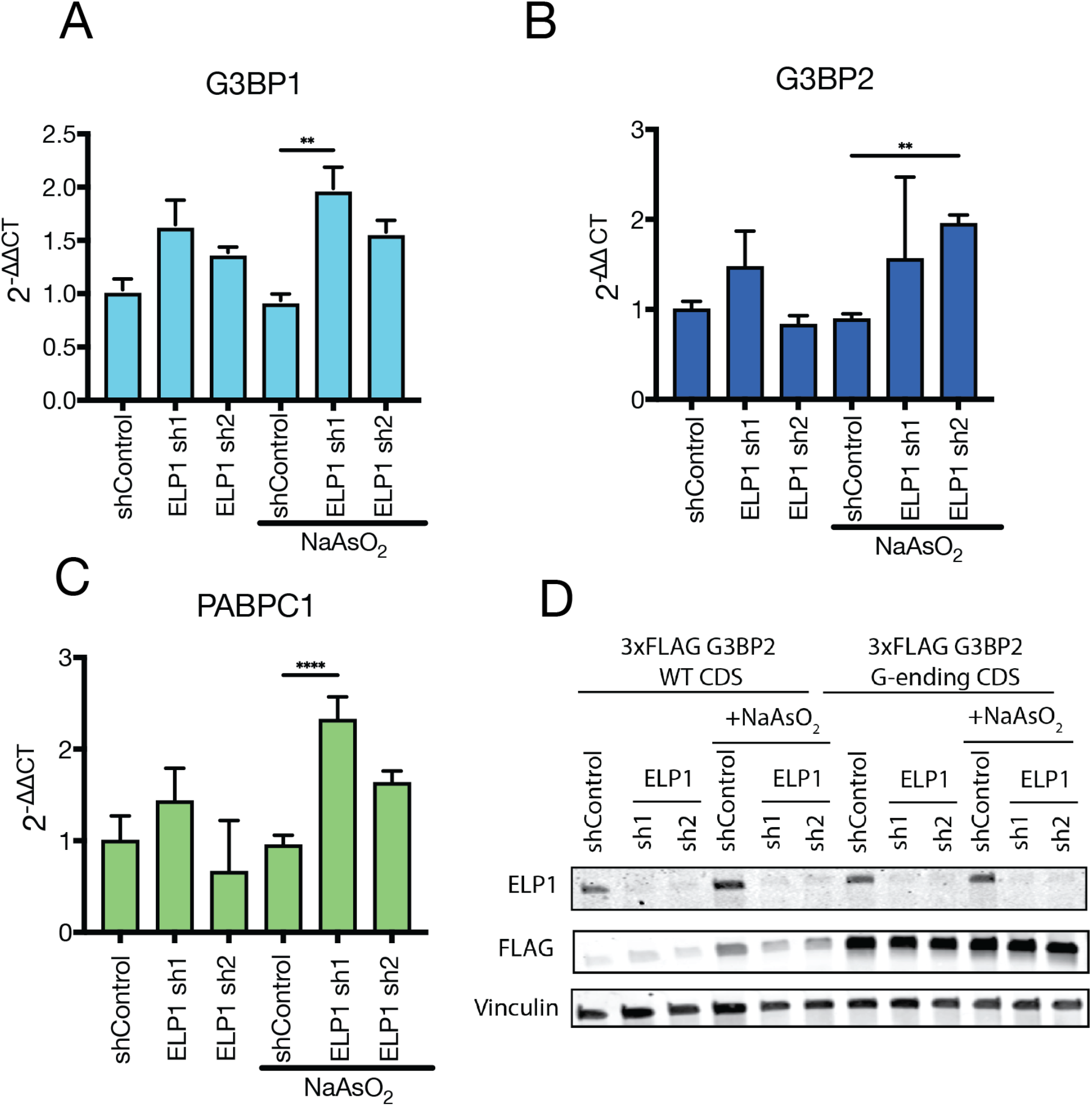
(A-C) Reverse transcriptase-qPCR of stress granule associated genes, (A) G2BP1, (B) G2BP2, and (C) PABPC1, in A375 melanoma treated with 100 μM NaAsO_2_ for 1 hr. (D) Western blot for of A375 melanoma cells treated with 100 μM NaAsO_2_ for 1 hr with overexpression of either wild type (WT) coding sequence (CDS) or G-ending codon switched CDS of G3BP2 upon loss of ELP1.

## Materials & Methods

### Reagents and Equipment

Commercially available reagents were used without further purification.

### Cell lines

Cells used for this study include A375 cells, A2058 cells, and HEK293T cells all purchased from ATCC. All cell lines were cultured in DMEM high glucose supplemented with 10% fetal bovine serum (Thomas Scientific) and 1x penicillin/streptomycin (Corning), unless otherwise specified.

### Plasmid Construction

miR-E shRNA knockdown constructs were generated by the splashRNA program and ordered from Invitrogen as oligos. Hairpins were ligated (T4 Ligase, NEB) into SGEN, SiREP, or SSEP lentiviral plasmid behind an SFFV promoter and EGFP, iRFP, or Scarlet fluorescent protein, respectively, using XhoI and EcoRI sites. Additionally, hairpins were ligated into UT4iREPIR, UT4GENIR, UT4SEPIR lentiviral plasmids behind a TRE-3GV dox inducible promoter, with either a iRFP, GFP, or Scarlet fluorescent protein and with either puro or neomycin resistance respectively. For overexpression constructs, the coding sequences of specific proteins were ordered as gene fragments (IDT) with an N-terminus kozack sequence. Reporter GFP fragemnts were ordederd as gBlocks with altered codon biases with all occurrences of mcm^5^s^2^U_34_ amino acids altered to the G-ending or A-ending version (AGG or AGA for Arg) (Integrated DNA Technologies) were ligated (In-Fusion HD, Takara) into a pLenti vector behind an EFS promoter using KflII and NsiI sites. An Internal ribosomal entry site (IRES) sequence links the EGFP reporter to an iRFP for normalization. Overexpression constructs of G3BP1, G3BP2, and PABPC1 were ordered as CDS fragments as either the WT CDS or with all occurrences of mcm^5^s^2^U_34_ amino acids altered to the G-ending version (AGG for Arg) from Twist Bioscience, these were ligated (In-Fusion HD, Takara) into pLenti vector behind an EFS promoter using BamHI and NsiI sites and sequence homology. Original vector backbones were kindly provided by Lukas Dow, and all plasmids were sequence verified.

### Lentiviral transduction and generation of stable cell lines

For virus production, 0.9μg of plasmid was combined with 1μg of packaging plasmids (0.4μg pMD2G and 0.6μg psPAX2) and transfected into HEK 293T cells using Polyjet (SignaGen) according to manufacturer’s instructions. Fresh media was added the following day, viral supernatants were collected 72hr after transfection, and filtered through a 0.45μM filter. Approximately 1 million cells were infected with viral supernatant and 10μg/mL polybrene (Sigma-Aldrich). Cells then underwent antibiotic selection with puromycin or hygromycin (Inviogen) depending on the antibiotic resistance of each plasmid.

### Doxycycline inducible hairpin treatment

Cells expressing UT4iREPIR, UT4GENIR, or UT4SEPIR were treated with 1 μg/mL of doxycycline treated media for 3 days so that experimental endpoints were collected on day 4 (24hr of experiment) or 5 (48 hr of experiment) depending on the experiment being run.

### tRNA biotin enrichment

Small RNA was extracted from cells or tissue using RNAzol RT (Molecular Research Center) according to manufacturer’s instructions. tRNA enrichment was performed as previously described (Songe-Moller et al., 2010). Briefly, M270 Streptavidin beads were washed then incubated with a biotin-labelled oligo specific to the 3’ end of the tRNA for 30min at RT with gentle shaking (LysUUU: CGCCCGAACAGGGACTTGAACCCTGGACCC-3Bio, IDT). Beads were then washed to remove excess oligo and resuspended in in 6xSSC buffer (Fisher Scientific). 40-60μg small RNA was also resuspended in 6xSSC buffer. The beads and RNA were incubated for 5 min separately at 75°C and then combined for an additional 5 min incubation. The combined beads and small RNA were then incubated for 40 min at RT with gentle shaking. Beads were then washed three times with 3x SSC, two times with 1x SSC, and 6 times with 0.1x SSC or until the absorbance at 260nm of the wash supernatant was 0. tRNA was eluted in 0.1x SSC by incubating the beads with buffer at 65°C for 3-4 min. tRNA was then precipitated by adding 1μL glycogen (Sigma-Aldrich) and 3M NaOAc to the tRNA elution followed by the addition of 4× volume 100% ethanol. Samples were then stored at -80°C overnight.

### Analysis of nucleoside modification by LCMS

Ethanol-precipitated tRNA was pelleted by centrifugation for 15min at 10,000rpm. After washing with 70% ethanol, the pellet was resuspended in 30μL H_2_O. tRNA was then boiled at 100°C for three minutes and then immediately placed in an ice water bath to quickly cool. Two units Nuclease P1 was added and digested for two hours at 45°C. 0.002 units of venom phosphodiesterase was added with 1/10 volume 1M ammonium bicarbonate and incubated for 2 hours at 37°C. 0.5 units alkaline phosphatase was then added and incubated for one hour. Once digestion was complete, the full volume was transferred to a 10kDa centrifugal filter to remove enzymes. Columns were centrifuged for 8 min at 14,000xg at 4°C. samples were stored at -80°C. Nucleoside samples were extracted using a solvent mixture of 46% methanol, 31% chloroform, 23% water followed by vortexing at 4 °C for 10 min at maximum speed. The extracts were centrifuged at 4 °C for 10 min at 21,300 × g in a microcentrifuge (Eppendorf Centrifuge 5425 R). The aqueous phase was transferred to 0.4 mL Polypropylene vials and dried down in SpeedVac concentrator (SPD120, ThermoFisher Scientific). Dried samples were resuspended in LC/MS-grade water prior to HPLC-MS analysis.

Nucleoside samples were separated using the Vanquish HPLC system (ThermoFisher Scientific) through an Atlantis Premier BEH Z-HILIC LC column (2.1 × 150 mm Column, 1.7 μm particle size) in a column oven temperature maintained at 30 °C. A 2 μL injection volume was used and an adjustable post-column follow splitter was set to a 1:1 ratio at a constant flow rate of 0.250 ml/min.

Buffer A consisted of 15mM ammonium formate (pH 5.5) in 30% acetonitrile and 70% water. Buffer B is 100% acetonitrile. The gradient profile was as follows: 86% buffer B (0.0-2.0 min), linear decrease from 86%-15% buffer B (2.0-14.0), 15% buffer B (14.0-16.0), linear increase from 15%-86% buffer B (16-16.4 min), and 86% buffer B (16.4-20 min).

Mass spectrometry data were acquired on an Orbitrap Exploris 120 mass spectrometer (ThermoFisher Scientific) equipped with a heated electrospray ionization (HESI) source. The method was a Top 4 method with a duration of 20 minutes. The spray voltage was set to 2.7 kV for positive and negative ion modes. The ion transfer tube temperature was 325 °C and the vaporizer temperature was 350 °C. Full MS scans were carried out at a resolution of 30,000 with a scan range of 67-1000 m/z in profile mode. Peak heights were measured using Xcalibur software (ThermoFisher Scientific) and corrected for natural abundance. Data was normalized to Cytosine signal and then normalized to control. For multiple group comparisons statistical analyses were performed using an ordinary two-way ANOVA followed by Dunnets multiple comparisons test, with a single pooled variance. Data are presented as mean ± standard deviation (SD), and a p-value < 0.05 was considered statistically significant.

### Flow cytometric analysis of OP-Puro incorporation

Following OP-Puro Kit (ThermoFisher), in brief cells in culture were plated at 30k cells/well in 24 well platesplates 24hrs prior to analysis. OP-Puro was added to the cells for 30 min at 37°C, final concentration 10μM. Cells were dissociated by trypsin (Corning). After OP-puro treatment, cells were fixed with Tonbo biosciences FOXP3 fix and perm kit. Cells were treated with the 647-azide and copper protectant (OP-Puro kit, ThermoFisher) for 30 min at Room temperature. Cells were resuspended in HBSS-free (Ca^2+^-and Mg^2+^-free, Gibco) with 0.5 μg/mL DAPI. Mean fluorescent intensity (MFI) of the 647-azide for cells that were positive for DAPI and SGEN (GFP) was observed.

For Melanoma cells from PDX tumors, mice were dissected rapidly for tumor sections and metastatic nodules. Both tumors and nodules were dissociated in enzymatic digestion media (see above) treated with OP-puro at final concentration of 20 μM for 30 min at 37°C shaking. The digests were resuspended in staining media (see above) and passed through a 100 μm cell strainer. After digestion and OP-puro treatment, cells were fixed with Tonbo biosciences FOXP3 fix and perm kit. Cells were treated with the 647-azide and copper protectant (OP-Puro kit, ThermoFisher) for 30 min at room temperature. Cells were then stained as described above for identification of melanoma cells and mean fluorescent intensity (MFI) of the 647-azide for cells positive for dsRed and HLA and negative for mouse CD45, CD31, Ter-119 was observed. Data was normalized to control. For multiple group comparisons statistical analyses were performed using an ordinary two-way ANOVA followed by Dunnets multiple comparisons test, with a single pooled variance. Data are presented as mean ± standard deviation (SD), and a p-value < 0.05 was considered statistically significant.

### Flow cytometric analysis of codon-biased GFP reporter for translational fidelity

Reporter constructs consisted of a destabilized GFP, with a half-life of 2 hrs, that was biased towards either the A-ending or G-ending codon by altering the coding sequence at each occurrence of the Lys, Glu, Gln, and Arg codons. An IRES and iRFP coding sequence immediately followed the GFP coding sequence. Cells stably expressing each individual reporter and the UT4SEPIR plasmid were plated 24hrs prior to analysis. Cells were dissociated by trypsin (corning) and resuspend in HBSS-free (Ca^2+^-and Mg^2+^-free, Gibco) with 0.5 μg/mL DAPI for analysis by Flow cytometry. GFP and Scarlet signals were compensated. Data was normalized to control. For multiple group comparisons statistical analyses were performed using an ordinary two-way ANOVA followed by Dunnets multiple comparisons test, with a single pooled variance. Data are presented as mean ± standard deviation (SD), and a p-value < 0.05 was considered statistically significant.

### Migration Assay

Plating into an Ibidi 4-well culture insert placed into a 12 well plate, 30,000 cells were added to each quadrant. After 24 hr, cells were treated with to mitomycin-C, final concentration 5 μg/mL, for 2 hrs. The inserts were removed, and the wells were gently washed twice with PBS and media was replaced. Cells were immediately imaged at the center and all four cardinal points of the insert’s cross for a time zero measurement. Cells were imaged every 24 hr until controls closed. 48 hr timepoint shown for A375 cells. Area closed was analyzed by ImageJ. For multiple group comparisons statistical analyses were performed using an ordinary two-way ANOVA followed by Dunnets multiple comparisons test, with a single pooled variance. Data are presented as mean ± standard deviation (SD), and a p-value < 0.05 was considered statistically significant.

### Invasion Assay

Corning Matrigel Invasion Chambers were thawed and rehydrated in plain media (no FBS) for 2 hr. Inserts were placed in 24 well plates that contained FBS media below the chamber. 100,000 cells were plated into Corning Matrigel Invasion Chambers in BSA media inside the chamber. Invasion was stopped at 24 or 48 hr by removing noninvaded cells from the inside of the chamber using a q-tip and PBS and fixing in the cells that invaded through the Matrigel with 100% ethanol for 15 min. The fixed cells were then stained with 0.2% Crytal Violet in 2% ethanol for 15 min and were imaged. Invasion was analyzed using a ImageJ plug-in. Coverage area was normalized to control. For multiple group comparisons statistical analyses were performed using an ordinary two-way ANOVA followed by Dunnets multiple comparisons test, with a single pooled variance. Data are presented as mean ± standard deviation (SD), and a p-value < 0.05 was considered statistically significant.

### Correlation of Codon-Biased Reporter Expression with Cell Migration

Stable A375 cell lines stably expressing A-ending or G-ending reporters, along with a Luc-P2A-GFP control line, were grown to confluence in 2D silicon wound-healing inserts (Ibidi). To inhibit proliferation, cells were incubated with 5 μM mitomycin C for 2 h, with 5 μM Hoechst 33342 added during the final 30 min of this treatment. Inserts were removed, cells were washed, and fresh DMEM (high glucose, no glutamine, no phenol red) supplemented with 10% FBS was added. Time-lapse images of four edge fields per cell line were captured every hour for 14 h on a Zeiss LSM 880 confocal microscope using a Plan-Apochromat 20×/0.8 M27 objective. Green (488 nm ex./516 nm em.), blue (405 nm ex./447 nm em.), and brightfield channels were acquired at 2048×2048 px resolution.

Raw image stacks were background-subtracted in Fiji-ImageJ using the rolling-ball algorithm with a 150-px radius.^42,43^ Cytoplasmic segmentation was performed in CellProfiler (v4.2.8)^44^, and fluorescence intensities of the A-and G-ending reporters and the Luc-P2A-GFP control were quantified. Migration trajectories of at least sixteen cells per field were traced manually using Fiji’s Manual Tracking plugin^45^, and total path lengths were extracted. Pearson’s correlation between reporter intensity and migration distance was calculated, with P values determined by a two-tailed test at 95% confidence. Linear regression lines were fit by the least-squares method.

### Patient derived xenografts

Melanoma specimens were obtained with informed consent from all patients according to protocls approved by the Institutional Review Boards of the University of Michigan Medical School (IRBMED approvals HUM00050754 and HUM00050085) and the University of Texas Southwestern Medical Center. Tumors were enzymatically digested in media containing 200 U/mL collagenase IV (Worthington) and 100 U/mL DNase for 30 min at 37 °C. Cells were filtered with a 40-100 μm cell strainer to obtain a single cell suspension. Specimens were frozen and sent to Weill Cornell Medicine under an MTA agreement.

### Lentiviral transduction of human melanoma cells

A lentiviral construct with luciferase and dsRed (luc P2A dsRed) was used to label patient-derived melanoma cells as described in Piskounova et al. Nature (2015). Virus was produced as described above. For lentiviral transduction, 750,000 patient derived melanoma cells freshly dissociated from primary tumors engrafted in mice were infected with viral supernatant and supplemented with 10 μg/mL polybrene (Sigma). The following day, the media was replaced and 48hrs post-infection cells were either injected subcutaneously into mice as bulk tumors or FACS sorted for positive infection. This was repeated for any genetic alterations for experiments using the shRNA plasmid SiREP.

### Cell labeling and sorting

All melanoma cells in this study stably express dsRed and luciferase so that melanoma cells could be distinguished by flow cytometry or bioluminescent imaging. When preparing cells for sorting by flow cytometry, cells were stained with antibodies against mouse CD45 (30-F11-VioletFluor, Tonbo), mouse CD31 (390-VioletFluor, eBiosciences), Ter119 (Ter-119-VioletFluor, Tonbo) and human HLA-A, -B, -C (BD Biosciences) to select live human melanoma cells and exclude mouse endothelial and hematopoietic cells. Antibody labelling was performed for 20 min on ice, followed by washing and centrifugation. Before sorting, cells were resuspended in staining medium (L15 medium containing bovine serum albumin (1 mg/mL), 1% penicillin/streptomycin, and 10mM HEPES, pH 7.4) containing 4’6-diamidino-2-phenylindole (DAPI; 5μg/mL; Sigma) to eliminate dead cells from sorting. Live human melanoma cells were isolated by flow cytometry by sorting cells that were positive for dsRed, iRFP, and HLA, and negative for mouse CD45, CD31, Ter-119 and DAPI.

### Animal subjects

6-10 week old male and female NSG (NOD scid gamma) immunodeficient mice were used for subcutaneous injections. NSG mice were bred by and obtained from Kvin Lertpiriyapong, Weill Cornell or JAX laboratories.

### Transplantation of melanoma cells

After sorting, cells were counted and resuspended in DMEM media with 50% high-protein Matrigel in L15 (product 354248, Corning). For a standard metastasis assay, 100 cells were injected subcutaneously into the right flank of the mice. Tumor formation was evaluated by regular palpitation of the injection site and tumors were measured weekly until any tumor in the mouse cohort reached 2.5cm in its largest diameter. Mice were monitored daily for signs of distress according to a standard body condition score or within 24hr of their tumors reaching 2.5 cm in diameter – whichever came first. These experiments were performed according to protocols approved by the Institutional Animal Care and Use Committee at Weill Cornell Medicine (protocol 2017-0033).

### Bioluminescence imaging

Mice injected subcutaneously with melanoma cells expressing luciferase were monitored until tumors reached 2.5cm in diameter. For bioluminescent imaging, mice were injected intraperitoneally with 100μL of DPBS containing D-luciferin monopotassium salt (40μg/mL, Goldbio) 5 min before imaging, followed by general anesthesia with isoflurane 2 min before imaging. IVIS Imaging System 200 Series (Caliper Life Sciences) with Living Image Software was used with the exposure time set to 10 seconds. After imaging the whole body, mice were euthanized and individual organs were dissected and imaged. The bioluminescent signal was quantified with ‘region of interest’ measurement tools in Living Image (Perkin Elmer) software. Data was normalized to control mean. For multiple group comparisons statistical analyses were performed using an ordinary two-way ANOVA followed by Dunnets multiple comparisons test, with a single pooled variance. Data are presented as mean ± standard deviation (SD), and a p-value < 0.05 was considered statistically significant. After imaging, tumors and organs were fixed in paraformaldehyde for histopathology.

### Ribosomal Profiling (Ribo-seq) Library Construction

The ribosomal-sequencing libraries were generated as previously described with the following modifications.^46^ Cells were snap-frozen, allowed to thaw on ice and lysed in lysis buffer (50 mM Tris-HCl (pH 7.4), 150 mM NaCl, 5 mM MgCl_2_, 1 mM DTT, 1% Triton X-100, 100 μg ml^−1^ cycloheximide, 100 μg ml^−1^ tigecycline and 15 U ml^−1^ DNaseI (Thermo Fisher Scientific, 89836)). RNA was quantified using a Qubit broad-range RNA kit (Thermo Fisher Scientific, Q10211). The ribosome footprint was generated using 15 U of RNase I (Biosearch Technologies, N6901K) to 300 μg of RNA in a reaction volume of 200 μl at room temperature for 45 min. Ribosomes were collected by ultracentrifugation, and the ribosome footprint was purified using TRIzol (Thermo Fisher Scientific, 15596018). RNA was subjected to size selection from 17 to 40 nucleotides in a 15% polyacrylamide TBE-urea gel and then dephosphorylated. The ribosome footprint was extracted from the gel in extraction buffer (50 mM Tris-HCl (pH 7.0), 300 mM NaCl, 1 mM EDTA and 0.25% SDS) and was subjected to rRNA depletion using a Human/Mouse/Rat riboPOOL kit (siTools Biotech sold by Galen Molecular in Noth America, dp-K024-000050) following the manufacturer’s instructions. The rRNA-depleted ribosome footprint was precipitated in 80% ethanol with glycogen at –20 °C overnight. The rRNA-depleted ribosome footprint was then ligated to a preadenylated linker (rApp-WWAGATCGGAAGAGCACACGTC). The footprint was reverse transcribed using SuperScript III reverse transcriptase (Thermo Fisher Scientific, 18080044) with a 5′-phosphorylated primer that contains 8-nucleotide unique molecular identifiers and two spacer residues (Phos-NNNNNNNNAGATCGGAAGAGCGTCGTGTA-iSp18-AA-iSp18-TAGACGTGTGCTC). cDNA was run on 10% nondenaturing gels to remove the nonligated linker and the reverse transcription primer. cDNA was then circularized using CircLigase I in the presence of 1 mM ATP. The library was PCR amplified using KAPA HiFi HotStart PCR Kit (Roche, 07958897001) and NEBNext Multiplex Oligos for Illumina Dual Index Primers Set 2 (New England Biolabs, E7780S). The library was purified twice using 1× AMPure XP Beads (BeckmanCoulter, A63881). The library was sequenced on an Illumina NextSeq 2000 (single-end, 100 cycles) or an Illumina NovaSeq 6000 (paired-end, 100 cycles). Protocol and guidance were kindly provided by Shino Murikami.

### RNA-seq processing for paired ribo-seq samples

Adapters were removed with cutadapt 3.5 (-a NNAGATCGGAAGAGCACACGTCTGAACTCCAGTCAC -A NNAGATCGGAAGAGCACACGTCTGAACTCCAGTCAC -u 8).

Subsequently RNA-seq reads were mapped to the human genome (Ensembl v90) using STAR 2.7.11a (--alignIntronMax 1 --outSAMmode NoQS --outSAMtype BAM SortedByCoordinate --alignEndsType Extend5pOfRead1 --outSAMattributes nM MD NH).

### Ribo-seq processing

Adapters were removed with cutadapt 3.5 (-a NNAGATCGGAAGAGCACACGTCTGAACTCCAGTCAC -u 8). Reads were mapped against human rRNA sequences (U13369.1) using bowtie 1.3.1 with standard parameters and all mapped reads were discarded. The same step was repeated with the genomes of Mycoplasma hominis, Mycoplasma hyorhinis, Mycoplasma fermentans and Acholeplasma laidlawii.

The remaining reads were mapped using STAR 2.7.11a to the human genome (Ensembl v90) with the same parameters as the RNA-seq data.

### Codon distribution relative to P-site position

The read count for the coding sequence of each isoform across all libraries was determined and the isoform with the maximal number of mapped reads for each gene was determined. The major isoform for each gene is defined as the isoform with the highest read coverage (number of reads / sequence length) with a total read count exceeding 90% of the max isoform.

The major isoforms were used as the basis to compute the codon distribution relative to the ribosome P-sites. The most likely P-site for each read was estimated and the P-site coverage for each codon over the coding sequence was computed. Then the average P-site coverage was computed for each sample. Finally, codon propensity over all samples can be computed as the P-site read count divided by the average P-site coverage for all codon-offset pairs (offset from -15 to +15 amino acids). Translation start and stop sites were excluded.

(code in https://github.com/erhard-lab/gedi/blob/main/Gedi/src/gedi/riboseq/analysis/CodonUsagePriceAnalysis.java)

### RNA/RIBO/TE differential analysis

Read counts were generated by the gedi pipeline (https://github.com/erhard-lab/gedi). Differential transcription, translation and translational efficiency analysis was done with deltaTE (10.1002/cpmb.108).

### Tissue RNA sequencing

RNA-sequencing libraries from primary and metastatic tumors from PDX models of melanoma metastasis were prepared by snap-freezing tissue in TRIzol. RNA was extracted according to manufacturer’s instructions using TRIzol (Thermo Fisher Scientific, 15596018) and quantified on a NanoDrop. RNA libraries were generated using TruSeq Stranded Total RNA Library Preparation (rRNA depletion and stranded RNA sequencing). The libraries were single-end sequenced for 100 cycles on an Illumina NextSeq 2000.

#### Analysis

Raw sequencing reads were demultiplexed between human and mouse using BBSplit from the BBMap suite v35.85^47^ to separate xenograft-derived transcripts from host contamination. Human reads were pseudo-aligned to a curated human transcriptome (MANE 1.3) using Kallisto v0.46.1^48^, with transcript-level abundances summarized to gene-level counts using tximport v1.3.0 ^49^. Differential expression analysis was performed using DESeq2 v1.42.0^50^ with a two-factor model to adjust for mouse-specific batch effects. Genes with adjusted p-values < 0.05 (Benjamini-Hochberg correction) were considered significantly differentially expressed between metastatic and primary tumors.

### Proteomic Analysis of PDX Melanoma Metastases and Matched Primary Tumors

#### Protein Extraction and digestion

Tissues were washed once in ice-cold PBS containing protease inhibitors to decrease the level of blood contamination and lysed by sonication in a probe sonicator (10 sec pulse @ 100% Amp) in 500 ul of the lysis buffer composed of 5% SDS 10 mM TCEP 20 mM CAA and 100 mM TRIS, pH=8. Lysates were incubated for 30 min @ 95 °C (1,200 rpm), centrifuged, and clear supernatants were used for subsequent processing.^51^

Proteins were precipitated on SP3 magnetic beads by 2-fold dilution with EtOH and washed 3 times with 200 μL of 85% EtOH to remove SDS traces. Proteins were digested with trypsin while on magnetic beads in 50 mM TRIS for four hours @ 37 °C (1,200 rpm).

Digests were acidified with 10% TFA to a final 0.5 % TFA, and the aliquots of 1 μg of peptide digests were loaded onto Evosep One C18 tips for subsequent analysis by LC-MS/MS.

#### LC-MS/MS

LC separation was performed online on EvosepOne LC utilizing Dr Maisch C18 AQ, 1.9μm beads (150μm ID, 15cm long, cat# EV-1106) analytical column. Peptides were gradient eluted from the column directly into Orbitrap HFX mass spectrometer using 88 min extended Evosep method (SPD15) at a flowrate of 220 nl/min. The mass spectrometer was operated in a data-independent acquisition mode (DIA)^52^ doing MS/MS fragmentation across 22 m/z windows after every full MS scan event (see table below).

| window | start_mz | end_mz | center | width |
| --- | --- | --- | --- | --- |
| 1 | 350 | 384 | 367 | 34 |
| 2 | 383 | 411 | 397 | 28 |
| 3 | 410 | 436 | 423 | 26 |
| 4 | 435 | 460 | 447.5 | 25 |
| 5 | 459 | 482 | 470.5 | 23 |
| 6 | 481 | 504 | 492.5 | 23 |
| 7 | 503 | 527 | 515 | 24 |
| 8 | 526 | 548 | 537 | 22 |
| 9 | 547 | 571 | 559 | 24 |
| 10 | 570 | 593 | 581.5 | 23 |
| 11 | 592 | 616 | 604 | 24 |
| 12 | 615 | 640 | 627.5 | 25 |
| 13 | 639 | 666 | 652.5 | 27 |
| 14 | 665 | 693 | 679 | 28 |
| 15 | 692 | 723 | 707.5 | 31 |
| 16 | 722 | 755 | 738.5 | 33 |
| 17 | 754 | 791 | 772.5 | 37 |
| 18 | 790 | 834 | 812 | 44 |
| 19 | 833 | 885 | 859 | 52 |
| 20 | 884 | 956 | 920 | 72 |
| 21 | 955 | 1071 | 1013 | 116 |
| 22 | 1070 | 1650 | 1360 | 580 |

High resolution full MS spectra were acquired with a resolution of 120,000, an AGC target of 3e6, with a maximum ion injection time of 60 ms, and scan range of 350 to 1650 m/z. Following each full MS scan 22 data-independent HCD MS/MS scans were acquired at the resolution of 30,000, AGC target of 3e6, stepped NCE of 22.5, 25 and 27.5.

#### Data Analysis

MS data were analyzed using Spectronaut software (https://biognosys.com/shop/spectronaut) and searched in directDIA mode against the *Homo sapiens* uniprot database (http://www.uniprot.org/). Database search was performed in integrated search engine Pulsar. For searching, the enzyme specificity was set to trypsin with the maximum number of missed cleavages set to 2. Oxidation of methionine was searched as variable modification; carbamidomethylation of cysteines was searched as a fixed modification. The false discovery rate (FDR) for peptide, protein, and site identification was set to 1%. Protein quantification was performed on the M2 level using the 3 most intense fragment ions per precursor.

Differential protein expression between metastatic and primary tumors was assessed using log2-transformed intensities following label-free quantification normalization. Log2 fold changes were calculated within individual mice to control for inter-animal variability. Statistical significance was determined by one-sample t-test on the individual fold changes with Benjamini-Hochberg correction for multiple comparisons (q < 0.001). Statistical analyses were performed in R version 4.3.2.^53^

### Codon Bias Analysis

Coding sequences (CDS) for all human transcripts were obtained from Ensembl Biomart (MANE 1.3). Relative synonymous codon usage (RSCU, Equation 1)^37^ values were calculated for each transcript using the seqinr v4.2-8 package.^54^

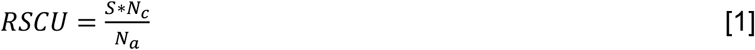

Where, N_c_= frequency of a specific codon in a gene

N_a_= Frequency of the amino acid N_c_ codes for in a gene

S=number of synonymous codons for N_c_

Used an RSCU value of greater than equivalent usage to classify biased proteins. In the case of Lys, Gln, and Glu, equivalent codon usage of synonymous codons is 50%; there are only two degenerate codons for these amino acids. For Arg, equivalent codon usage between all synonymous codons is 16.67%, as there are 6 degenerate codons, with only one requiring mcm^5^s^2^U_34_ modified tRNA for decoding.

### Stress Granule Induction

Cells were plated for the desired experiment. Cells were treated with sodium (meta)arsenite (NaAsO_2_, Sigma) treated media at 100 μM for 1hr at 37°C.

### Immunofluorescence Staining of Melanoma Cells

Cells were plated on glass slides and treated as described in the text. Cells were then fixed with 4% paraformaldehyde for 20 minutes at room temperature. Slides are washed 3 times with PBS followed with permeabilization either in PBS with 0.1% Triton X-100 (G3BP1) or methanol (PABP1) (at -20C). Cells are washed with PBS. Cells were blocked with 5% bovine serum albumin in TBST for 20 min at room temperature. Cells were then stained with the primary antibodies overnight at 4 °C in 5% bovine serum albumin in TBST. The next day, sections were washed in TBST twice for 5 min. Cells are stained with secondary antibody, goat anti-rabbit (A-21244, Invitrogen; 1:500), for 30 min in the dark at room temperature. Slides were washed with TBST twice. Cells were stained with DAPI (1:2000 in HBSS) for 2 minutes at room temperature. Cells were washed twice with TBST and once with diH2O and mounted for imaging. Fluorescent signal was analyzed and quantified using QuPath v0.4.3 software. For multiple group comparisons statistical analyses were performed using an ordinary two-way ANOVA followed by Dunnets multiple comparisons test, with a single pooled variance. Data are presented as mean ± standard deviation (SD), and a p-value < 0.05 was considered statistically significant.

### Immunohistochemistry of tissue sections

Tissues were fixed in 4% paraformaldehyde for 12 h at 4 °C. Fixed organs were embedded in paraffin and sectioned by IDEX, all organs and primary tumor were placed in one block. Sections (10 μm) were deparaffinized: xylene 5 min, xylene 5 min, 100% isopropanol 5 min, 100% isopropanol 5 min, 96% ethanol 2 min, 90% ethanol 3 min, 70% ethanol 2 min, dH2O 10 min. Slides were then subjected to antigen retrieval for 30 min boiling in 0.1M sodium citrate, and cooled to RT. Sections were blocked in 1x PGB (5mg/mL BSA, 1mg/mL Fish gelatine in 1X PBS) for 1 hr. Tissues are stained with primary antibody overnight at 4 °C in 1xPGB. Tissues were washed three times in 1X PBS for 5 min each time. After washing, sections were incubated with fluorophore-conjugated secondary antibodies for 1 hour. Nuclei were visualized with DAPI (4′,6-diamidino-2-phenylindole). Imaging was performed using a FluoView FV-10i Olympus laser scanning confocal microscope with a 60× objective. Excitation/emission wavelengths used were: 490/525 nm (Alexa Fluor 488) and 358/461 (DAPI). Image analysis was conducted using ImageJ software (version 1.53k). All statistical analyses were performed using GraphPad Prism version 10.5.0 (GraphPad Software). For multiple group comparisons statistical analyses were performed using an ordinary one-way ANOVA followed by Dunnets multiple comparisons test, with a single pooled variance. Data are presented as mean ± standard deviation (SD), and a p-value < 0.05 was considered statistically significant.

### Real-Time PCR Quantification of tRNA and mRNA

mRNA was extracted using a classic Trizol-chloroform RNA extraction. Reverse transcription was performed using iScript cDNA synthesis kit (Bio-Rad) as described by manufacturer. cDNA was then diluted and used for qPCR analysis with SYBR Green PCR master mix (Thermo Fisher). IDT online primer design tool was used to generate qPCR primers and UCSC In-Silico PCR database was then used to verify human-specificity of primers. qPCR analysis was performed on an Applied Biosystems QuantStudio 6 Real-Time PCR System. All targets were normalized to Actin in a standard ΔΔCt analysis.

### Statistical analysis

No statistical methods were used to predetermine sample size. The data in most figure panels represents several independent experiments performed on different days. Variation is always indicated using standard deviation. For analysis of statistical significance, we first tested whether there was homogeneity of variation across conditions (as required by ANOVA) using Levene’s test, or when only two conditions were compared, using the F-test. In cases where the variation was significantly different among conditions, we used a non-parametric Kruskal-Wallis test or a non-parametric Mann-Whitney test to assess significance of difference among populations and conditions. Usually, variation did not significantly differ among conditions. Under those circumstances, two-tailed Student’s t-tests were used to test significance of differences between two conditions. When more than one conditions were compared, a one-way ANOVA followed by Dunnett’s multiple comparisons test was performed. A two-way ANOVA followed by Dunnett’s multiple comparisons test were used in cases where more than two groups were compared with repeated measures. In all xenograft assays, we injected 5-8 week old NSG mice, 5 per condition. Both male and female mice were used. When mice died before the end of the experiment due to opportunistic infections the data from those mice was excluded.

## References

1. Dedon, P.C., and Begley, T.J. (2022). Dysfunctional tRNA reprogramming and codon-biased translation in cancer. Trends Mol Med 28, 964–978. 10.1016/j.molmed.2022.09.007.

2. Suzuki, T. (2021). The expanding world of tRNA modifications and their disease relevance. Nat. Rev. Mol. Cell Biol. 22, 375–392. 10.1038/s41580-021-00342-0.

3. Fernandez-Vazquez, J., Vargas-Perez, I., Sanso, M., Buhne, K., Carmona, M., Paulo, E., Hermand, D., Rodriguez-Gabriel, M., Ayte, J., Leidel, S., and Hidalgo, E. (2013). Modification of tRNA(Lys) UUU by elongator is essential for efficient translation of stress mRNAs. PLoS Genet 9, e1003647. 10.1371/journal.pgen.1003647.

4. Nedialkova, D.D., and Leidel, S.A. (2015). Optimization of Codon Translation Rates via tRNA Modifications Maintains Proteome Integrity. Cell 161, 1606–1618. 10.1016/j.cell.2015.05.022.

5. Delaunay, S., Rapino, F., Tharun, L., Zhou, Z., Heukamp, L., Termathe, M., Shostak, K., Klevernic, I., Florin, A., Desmecht, H., et al. (2016). Elp3 links tRNA modification to IRES-dependent translation of LEF1 to sustain metastasis in breast cancer. J Exp Med 213, 2503–2523. 10.1084/jem.20160397.

6. Rapino, F., Delaunay, S., Rambow, F., Zhou, Z., Tharun, L., De Tullio, P., Sin, O., Shostak, K., Schmitz, S., Piepers, J., et al. (2018). Codon-specific translation reprogramming promotes resistance to targeted therapy. Nature 558, 605–609. 10.1038/s41586-018-0243-7.

7. Rapino, F., Zhou, Z., Sanchez, A.M.R., Joiret, M., Seca, C., Hachem, N.E., Valenti, G., Latini, S., Shostak, K., Geris, L., et al. (2021). Wobble tRNA modification and hydrophilic amino acid patterns dictate protein fate. Nature Communications 12, 2170. 10.1038/s41467-021-22254-5.

8. Nedialkova, Danny D., and Leidel, Sebastian A. (2015). Optimization of Codon Translation Rates via tRNA Modifications Maintains Proteome Integrity. Cell 161, 1606–1618. 10.1016/j.cell.2015.05.022.

9. Chujo, T., and Tomizawa, K. (2021). Human transfer RNA modopathies: diseases caused by aberrations in transfer RNA modifications. FEBS J. 288, 7096–7122. 10.1111/febs.15736.

10. Crick, F.H. (1966). Codon--anticodon pairing: the wobble hypothesis. J Mol Biol 19, 548–555. 10.1016/s0022-2836(66)80022-0.

11. El Yacoubi, B., Bailly, M., and de Crecy-Lagard, V. (2012). Biosynthesis and function of posttranscriptional modifications of transfer RNAs. Annu Rev Genet 46, 69–95. 10.1146/annurev-genet-110711-155641.

12. Linder, B., Sharma, P., Wu, J., Birbaumer, T., Eggers, C., Murakami, S., Ott, R.E., Fenzl, K., Vorgerd, H., Erhard, F., et al. (2025). tRNA modifications tune m(6)A-dependent mRNA decay. Cell. 10.1016/j.cell.2025.04.013.

13. Murakami, S., Olarerin-George, A.O., Liu, J.F., Zaccara, S., Hawley, B., and Jaffrey, S.R. (2025). m(6)A alters ribosome dynamics to initiate mRNA degradation. Cell. 10.1016/j.cell.2025.04.020.

14. Hughes, R.O., Davis, H.J., Nease, L.A., and Piskounova, E. (2024). Decoding the role of tRNA modifications in cancer progression. Curr Opin Genet Dev 88, 102238. 10.1016/j.gde.2024.102238.

15. Piskounova, E., Agathocleous, M., Murphy, M.M., Hu, Z., Huddlestun, S.E., Zhao, Z., Leitch, A.M., Johnson, T.M., DeBerardinis, R.J., and Morrison, S.J. (2015). Oxidative stress inhibits distant metastasis by human melanoma cells. Nature 527, 186–191. 10.1038/nature15726.

16. Ganesh, K., and Massague, J. (2021). Targeting metastatic cancer. Nat Med 27, 34–44. 10.1038/s41591-020-01195-4.

17. Arfin, S., Jha, N.K., Jha, S.K., Kesari, K.K., Ruokolainen, J., Roychoudhury, S., Rathi, B., and Kumar, D. (2021). Oxidative Stress in Cancer Cell Metabolism. Antioxidants (Basel) 10. 10.3390/antiox10050642.

18. Lu, J., Tan, M., and Cai, Q. (2015). The Warburg effect in tumor progression: mitochondrial oxidative metabolism as an anti-metastasis mechanism. Cancer Lett 356, 156–164. 10.1016/j.canlet.2014.04.001.

19. Ramundo, V., Giribaldi, G., and Aldieri, E. (2021). Transforming Growth Factor-beta and Oxidative Stress in Cancer: A Crosstalk in Driving Tumor Transformation. Cancers (Basel) 13. 10.3390/cancers13123093.

20. Lu, J.J., Abudukeyoumu, A., Zhang, X., Liu, L.B., Li, M.Q., and Xie, F. (2021). Heme oxygenase 1: a novel oncogene in multiple gynecological cancers. Int J Biol Sci 17, 2252–2261. 10.7150/ijbs.61073.

21. Cascio, G., Aguirre, K.N., Church, K.P., Hughes, R.O., Nease, L.A., Delclaux, I., Davis, H.J., and Piskounova, E. (2024). Transcriptional Isoforms of NAD(+) kinase regulate oxidative stress resistance and melanoma metastasis. Redox Biol 76, 103289. 10.1016/j.redox.2024.103289.

22. Leona A. Nease, K.P.C., Ines Delclaux, Shino Murakami, Maider Astorkia, Marwa Zerhouni, Graciela Cascio, Riley O. Hughes, Kelsey N. Aguirre, Paul Zumbo, Lukas E. Dow, Samie Jaffrey, Doron Betel & Elena Piskounova (2024). Selenocysteine tRNA methylation promotes oxidative stress resistance in melanoma metastasis. Nat. Cancer. 10.1038/s43018-024-00844-8.

23. Ganesh, K., and Massagué, J. (2021). Targeting metastatic cancer. Nature Medicine 27, 34–44. 10.1038/s41591-020-01195-4.

24. Reiter, J.G., Baretti, M., Gerold, J.M., Makohon-Moore, A.P., Daud, A., Iacobuzio-Donahue, C.A., Azad, N.S., Kinzler, K.W., Nowak, M.A., and Vogelstein, B. (2019). An analysis of genetic heterogeneity in untreated cancers. Nat Rev Cancer 19, 639–650. 10.1038/s41568-019-0185-x.

25. Zhou, H., Luo, J., Mou, K., Peng, L., Li, X., Lei, Y., Wang, J., Lin, S., Luo, Y., and Xiang, L. (2023). Stress granules: functions and mechanisms in cancer. Cell Biosci 13, 86. 10.1186/s13578-023-01030-6.

26. Youn, J.Y., Dyakov, B.J.A., Zhang, J., Knight, J.D.R., Vernon, R.M., Forman-Kay, J.D., and Gingras, A.C. (2019). Properties of Stress Granule and P-Body Proteomes. Mol Cell 76, 286–294. 10.1016/j.molcel.2019.09.014.

27. Ge, Y., Jin, J., Li, J., Ye, M., and Jin, X. (2022). The roles of G3BP1 in human diseases (review). Gene 821, 146294. 10.1016/j.gene.2022.146294.

28. Ryan, L., and Rubinsztein, D.C. (2024). The autophagy of stress granules. FEBS Lett 598, 59–72. 10.1002/1873-3468.14787.

29. Wheeler, J.R., Matheny, T., Jain, S., Abrisch, R., and Parker, R. (2016). Distinct stages in stress granule assembly and disassembly. Elife 5. 10.7554/eLife.18413.

30. Matsuki, H., Takahashi, M., Higuchi, M., Makokha, G.N., Oie, M., and Fujii, M. (2013). Both G3BP1 and G3BP2 contribute to stress granule formation. Genes Cells 18, 135–146. 10.1111/gtc.12023.

31. Tourrière, H., Chebli, K., Zekri, L., Courselaud, B., Blanchard, J.M., Bertrand, E., and Tazi, J. (2023). The RasGAP-associated endoribonuclease G3BP mediates stress granule assembly. J. Cell Biol. 222, e200212128072023new. 10.1083/jcb.200212128072023new.

32. Yang, P., Mathieu, C., Kolaitis, R.-M., Zhang, P., Messing, J., Yurtsever, U., Yang, Z., Wu, J., Li, Y., Pan, Q., et al. (2020). G3BP1 Is a Tunable Switch that Triggers Phase Separation to Assemble Stress Granules. Cell 181, 325-345.e328. 10.1016/j.cell.2020.03.046.

33. Grabocka, E., and Bar-Sagi, D. (2016). Mutant KRAS Enhances Tumor Cell Fitness by Upregulating Stress Granules. Cell 167, 1803–1813 e1812. 10.1016/j.cell.2016.11.035.

34. Abbassi, N.-e.-H., Jaciuk, M., Scherf, D., Böhnert, P., Rau, A., Hammermeister, A., Rawski, M., Indyka, P., Wazny, G., Chramiec-Głąbik, A., et al. (2024). Cryo-EM structures of the human Elongator complex at work. Nature Communications 15, 4094. 10.1038/s41467-024-48251-y.

35. Banh, R.S., Biancur, D.E., Yamamoto, K., Sohn, A.S.W., Walters, B., Kuljanin, M., Gikandi, A., Wang, H., Mancias, J.D., Schneider, R.J., et al. (2020). Neurons Release Serine to Support mRNA Translation in Pancreatic Cancer. Cell 183, 1202–1218 e1225. 10.1016/j.cell.2020.10.016.

36. Chothani, S., Adami, E., Ouyang, J.F., Viswanathan, S., Hubner, N., Cook, S.A., Schafer, S., and Rackham, O.J.L. (2019). deltaTE: Detection of Translationally Regulated Genes by Integrative Analysis of Ribo-seq and RNA-seq Data. Curr Protoc Mol Biol 129, e108. 10.1002/cpmb.108.

37. Xu, X.Z., Liu, Q.P., Fan, L.J., Cui, X.F., and Zhou, X.P. (2008). Analysis of synonymous codon usage and evolution of begomoviruses. J Zhejiang Univ Sci B 9, 667–674. 10.1631/jzus.B0820005.

38. Riggs, C.L., Kedersha, N., Ivanov, P., and Anderson, P. (2020). Mammalian stress granules and P bodies at a glance. J. Cell Sci. 133, jcs242487. 10.1242/jcs.242487.

39. Schwanhausser, B., Busse, D., Li, N., Dittmar, G., Schuchhardt, J., Wolf, J., Chen, W., and Selbach, M. (2011). Global quantification of mammalian gene expression control. Nature 473, 337–342. 10.1038/nature10098.

40. Vogel, C. (2011). Translation’s coming of age. Mol Syst Biol 7, 498. 10.1038/msb.2011.33.

41. Libert, M., Quiquempoix, S., Fain, J.S., Ruys, S.P.d., Haidar, M., Wulleman, M., Herinckx, G., Vertommen, D., Bouchart, C., Arsenijevic, T., et al. (2024). Stress granules are not present in Kras mutant cancers and do not control tumor growth. EMBO Rep. 25, 4693–4707. 10.1038/s44319-024-00284-6.

42. Schindelin, J., Arganda-Carreras, I., Frise, E., Kaynig, V., Longair, M., Pietzsch, T., Preibisch, S., Rueden, C., Saalfeld, S., Schmid, B., et al. (2012). Fiji: an open-source platform for biological-image analysis. Nat Methods 9, 676–682. 10.1038/nmeth.2019.

43. Sternberg, S.R. (1983). Biomedical Image-Processing. Computer 16, 22–34.

44. Stirling, D.R., Swain-Bowden, M.J., Lucas, A.M., Carpenter, A.E., Cimini, B.A., and Goodman, A. (2021). CellProfiler 4: improvements in speed, utility and usability. BMC Bioinformatics 22, 433. 10.1186/s12859-021-04344-9.

45. Cordelières, F. Manual Tracking plugin for ImageJ/Fiji.

46. McGlincy, N.J., and Ingolia, N.T. (2017). Transcriptome-wide measurement of translation by ribosome profiling. Methods 126, 112–129. 10.1016/j.ymeth.2017.05.028.

47. Bushnell, B. (2014). BBMap: a fast, accurate, splice-aware aligner (Lawrence Berkeley National Laboratory).

48. Bray, N.L., Pimentel, H., Melsted, P., and Pachter, L. (2016). Near-optimal probabilistic RNA-seq quantification. Nat Biotechnol 34, 525–527. 10.1038/nbt.3519.

49. Soneson, C., Love, M.I., and Robinson, M.D. (2015). Differential analyses for RNA-seq: transcript-level estimates improve gene-level inferences. F1000Res 4, 1521. 10.12688/f1000research.7563.2.

50. Love, M.I., Huber, W., and Anders, S. (2014). Moderated estimation of fold change and dispersion for RNA-seq data with DESeq2. Genome Biol 15, 550. 10.1186/s13059-014-0550-8.

51. Doellinger, J., Schneider, A., Hoeller, M., and Lasch, P. (2020). Sample Preparation by Easy Extraction and Digestion (SPEED) - A Universal, Rapid, and Detergent-free Protocol for Proteomics Based on Acid Extraction. Mol Cell Proteomics 19, 209–222. 10.1074/mcp.TIR119.001616.

52. Ludwig, C., Gillet, L., Rosenberger, G., Amon, S., Collins, B.C., and Aebersold, R. (2018). Data-independent acquisition-based SWATH-MS for quantitative proteomics: a tutorial. Mol Syst Biol 14, e8126. 10.15252/msb.20178126.

53. Team, R.C. (2023). R: A Language and Environment for Statistical Computing (R Foundation for Statistical Computing, Vienna, Austria).

54. Charif, D.a.L., J.R (2007). SeqinR 1.0-2: a contributed package to the R project for statistical computing devoted to biological sequences retrieval and analysis. In Structural approaches to sequence evolution: Molecules, networks, populations. Biological and Medical Physics, Biomedical Engineering series, U. Bastolla, Porto, M., Roman, H.E., and Vendruscolo, M., ed. (Springer Verlag), pp. pp. 207–232.

